# Procrustes Alignment in Individual-level Analyses of Functional Gradients

**DOI:** 10.1101/2024.11.26.625368

**Authors:** Leonard Sasse, Casey Paquola, Juergen Dukart, Felix Hoffstaedter, Simon B. Eickhoff, Kaustubh R. Patil

## Abstract

Functional connectivity (FC) gradients provide valuable insights into individual differences in brain organization, yet aligning these gradients across individuals poses challenges. Procrustes alignment is often employed to standardize gradients across multiple subjects, but the choice of the number of gradients used in alignment introduces complexities that may impact individual-level analyses. In this study, we systematically investigate the impact of varying gradient counts in Procrustes alignment on the principal FC gradient, using data from four resting state fMRI datasets, including the Human Connectome Project (HCP-YA), Amsterdam Open MRI Collection (AOMIC) PIOP1 and PIOP2, and Cambridge Centre for Ageing and Neuroscience (Cam-CAN). We find that increasing the number of gradients used in alignment enhances identification accuracy but can reduce differential identifiability, as additional gradients risk introducing nuisance signals such as motion back into the principal gradient. To further probe these effects, machine learning to predict fluid intelligence and age, and a motion prediction analysis, revealing that higher alignment gradient counts may leak information from lower gradients into the principal gradient for individual-level analyses. These findings highlight the trade-off between alignment precision and the potential reintroduction of noise.

**Key Points:**

- Gradient count used in Procrustes alignment impacts identification accuracy and differential identifiability obtained using the principal gradient
- The magnitude of the Procrustes transformation correlates with motion measures, and this correlation increases with higher gradient count used in alignment.
- Gradient count used in Procrustes alignment impacts prediction of fluid intelligence and age

## 1. Introduction

Functional connectivity (FC) derived using functional magnetic resonance imaging (fMRI) is a cornerstone of research for unravelling human brain organisation (Biswal et al., 1995; Craddock et al., 2013; Zalesky et al., 2012). By examining the temporal relationships between different brain regions, FC analysis has helped unveil intricate networks that underlie various cognitive processes and behaviours (Fox et al., 2005; Yeo et al., 2011). FC is also commonly used for investigating individual differences in brain-behaviour associations (Amico & Goñi, 2018; Chen et al., 2021; Finn et al., 2015; He et al., 2020; Sasse et al., 2023; Shen et al., 2017). Beyond traditional FC analysis, more abstract representations such as functional gradients have gained popularity (Bijsterbosch et al., 2020; 2021; Haak et al., 2018) for their ability to reveal systematic patterns of connectivity across the brain (Margulies et al., 2016; Wael et al., 2020). The “principal gradient” represents the primary axis of variance from unimodal sensorimotor cortical areas towards heteromodal association cortices, thereby resembling the hierarchical organisation of the cortex identified in prior anatomical work (Margulies et al., 2016; Mesulam, 1998). Notably, gradients have shown some potential in the study of inter-individual differences as a reliable and predictive feature (Alberti et al., 2023; Gonzalez Alam et al., 2022; Hong et al., 2020; Knodt et al., 2023; Kong et al., 2023). For instance, individual differences in functional gradients have been related to variables of clinical interest such as age or autism diagnosis (Bethlehem et al., 2017; 2020; Wan et al., 2023).

When gradients are computed from functional connectivity matrices, each parcel (i.e. distinct region of the brain) is assigned a value along the gradient. These values can vary in sign across individuals, meaning that the direction of the gradient can be flipped. Additionally, gradients are ordered based on their eigenvalues, but this ordering of gradients can also differ between individuals. Therefore, the gradients need to be aligned to make them comparable. Procrustes alignment is often employed for this purpose (Coifman & Hirn, 2014; Hong et al., 2020; Wael et al., 2020). Procrustes alignment aims to bring the gradients from different individuals into a common space using a group-level gradient as a reference, thereby simplifying group-level analyses (Bethlehem et al., 2020; Kim et al., 2024).

When performing Procrustes alignment one can choose the number of gradients to align simultaneously. It has been recommended to use 10 gradients for alignment, in order to maximise the fit between the group- and individual-level gradients (Hardikar et al., 2024; Mckeown et al., 2020) which is also the default in the most commonly used gradients toolbox, BrainSpace (version 0.1.10, (Wael et al., 2020)). However, it is common to restrict the downstream analysis only to the first few gradients, for instance three, arguably because some biological interpretation can be attributed to them (Margulies et al., 2016; Zhang et al., 2019). Given Procrustes alignment transforms each gradient based on information from all the gradients used in the alignment procedure, the number of gradients used for alignment can considerably impact the outcome. Therefore, understanding the impact of this choice in empirical research is crucial. Further, if Procrustes alignment does use information obtained from all the gradients to transform a specific gradient (e.g. the principal gradient is transformed using all 10 gradients used in alignment), it is critical to understand the origin and nature of the signal used in this process. This is especially critical if the transformation captures noise or relies on unwanted signals. For example, motion signals can persist in FC data even after extensive processing independent of the specific denoising pipeline chosen (Ciric et al., 2017; Kopal et al., 2020; Power et al., 2012; Satterthwaite et al., 2019; Siegel et al., 2017).

Individual-level analysis is a promising and highly anticipated application of FC and, by extension, of FC gradients. However, the impact of Procrustes alignment on individual-level analysis such as machine-learning-based prediction has not been fully accessed. As different individuals need alignment to a different extent, this consideration becomes crucial when investigating associations of specific gradients with individual-level behaviour, cognition and disease status. It is possible that individual characteristics might affect the alignment, and if so, then which individual characteristics contribute to the extent of the required transformations has not been explored. Specifically, using a higher number of gradients in the alignment process allows for a greater capacity to fit the data. However, this also increases the risk that nuisance signals, such as residual motion artefacts or noise from other unwanted sources, may get incorporated into the principal gradient during the alignment process, because Procrustes alignment transforms each gradient based on the information derived from all gradients used in the alignment. When more gradients are included, there is a higher likelihood that these additional gradients capture not just meaningful neural signals but also unwanted variability. As a result, the principal gradient - which is often of primary interest due to its biological interpretability - could become contaminated with signals that researchers intended to eliminate during the preprocessing stages. Clarifying those questions is essential for establishing gradients as a biomarker for individual-level analysis.

Here, we investigate the impact of the number of gradients used in Procrustes alignment on the aligned principal gradient, and its use in subject-level downstream analyses across different datasets including the high quality resting fMRI data available in the Human Connectome Project (HCP-YA) S1200 dataset (Van Essen et al., 2013), as well as resting state fMRI data from the Amsterdam Open MRI Collection (AOMIC) (Snoek et al., 2021) and the Cambridge Centre for Ageing and Neuroscience (Cam-CAN) (Taylor et al., 2017). In particular, we test for the impact of motion on the magnitude of the Procrustes transformation and its relationship to the number of gradients used in alignment. In addition, we also investigate the impact of Procrustes alignment on the principal gradients capacity to predict fluid intelligence and age using a machine learning approach.

## 2. Results

### 2.1. Identification and differential identifiability

First, we explored the impact of the number of gradients used in Procrustes alignment on subject-level analyses by evaluating two key metrics: identification accuracy and differential identifiability (see Section 4.4). These metrics provide insight into how well the aligned principal gradient can capture individual-level differences. Identification accuracy refers to the proportion of participants who were matched with highest correlation between two fMRI sessions based on their principal gradient. Differential identifiability measures the difference between the mean within-subject correlations and the mean between-subject correlations. However, Pearson’s r is not a proper distance metric. For instance, Pearson’s r is not strictly equidistant (see Section 4.4) (Solo, 2019). Therefore we performed an additional analysis in which Fisher’s r-to-z transform is first applied to all correlation values before computing differential identifiability. Importantly, in all our downstream analyses, we only use the first (“principal”) gradient, idenpendent of how many gradients were used in Procrustes alignment.

Identification accuracy using the principal gradient increased substantially and monotonically with the number of gradients used in Procrustes alignment (Fig. 1a), whereas the differential identifiability without Fisher’s r-to-z transform decreased (Fig. 1b). This result may seem counterintuitive at first glance because for both metrics higher values mean better identification. However, this can happen if both within- and between-subject correlation increases but the latter does not increase enough to cause misidentification (see supplementary; Fig. S1). In addition, however, the lack of equidistance in a metric like Pearson’s correlation could cause differential identifiability to decrease because inconsistent spacing between correlation values may distort the difference between within- and between-subject similarity measures, making it appear as though subjects are less distinguishable, even when their individual patterns become more unique. And indeed, after applying Fisher’s r-to-z transform to correlation values before calculating differential identifiability, we find that differential identifiability behaves more consistently with identification accuracy (Fig. 1c). In other words, differential identifiability with Fisher’s r-to-z transform applied to within- and between-subject correlations also increases with increasing number of gradients used in alignment. Since these findings were consistent across all dimensionality reduction approaches, in subsequent analyses, we focused on gradient extraction using the following recommended parameters: kernel = normalized_angle, sparsity = 0.9, dimensionality reduction technique = diffusion map embedding (“dm”) (see Section 4.2; (Bethlehem et al., 2017; 2020; Hänisch et al., 2023; Kong et al., 2023; Wan et al., 2023)).

**Figure 1:**
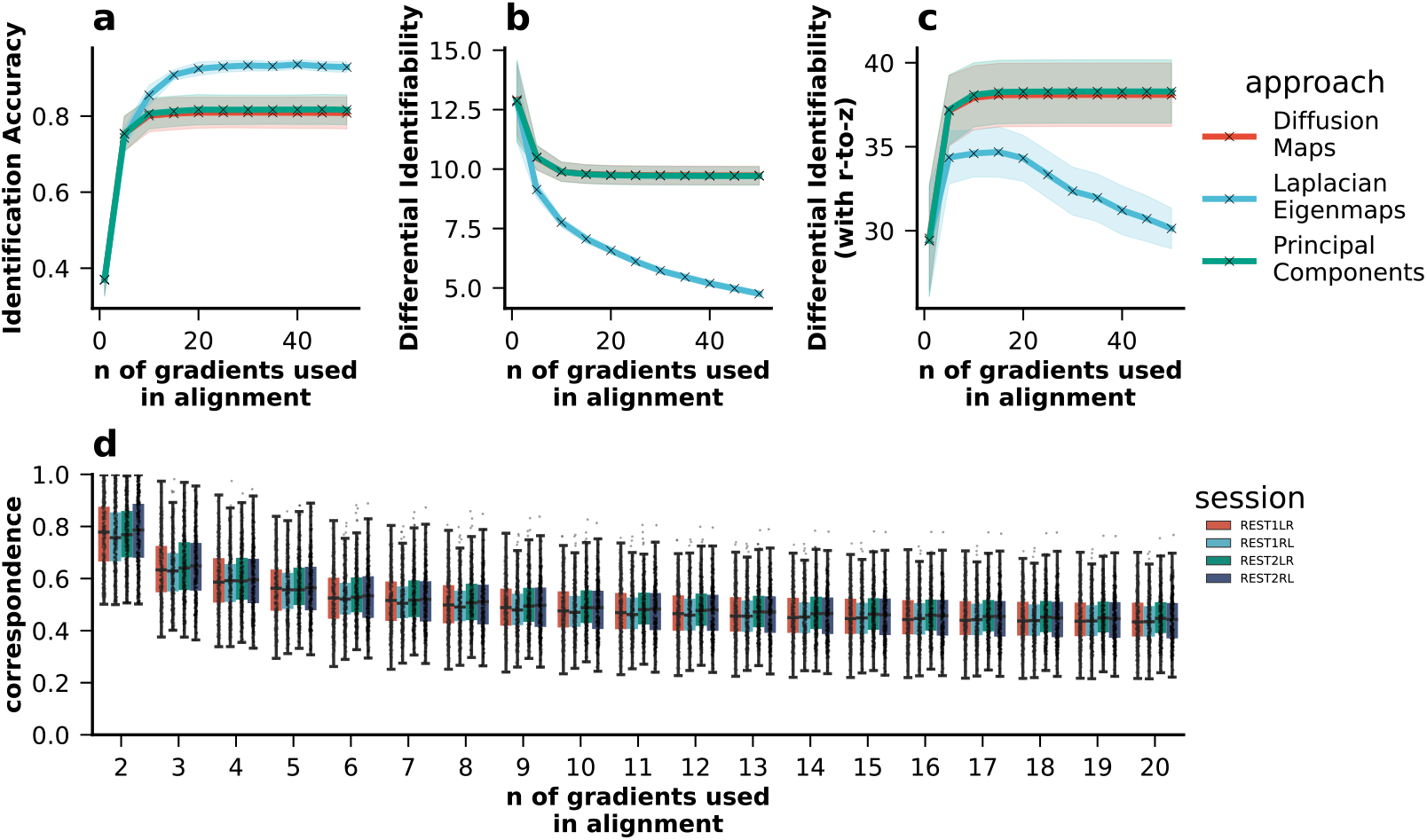
Impact of Procrustes alignment on a) identification accuracy and b) differential identifiability. For each subject, gradients were extracted per session (kernel = normalized_angle; sparsity = 0.9). They were then aligned to the holdout reference gradient using Procrustes alignment. Identification accuracy and differential identifiability were calculated for each combination of sessions (N_Sessions_ = 4; N_Combinations_ = 6). c) Lastly, the correspondence between the unaligned and the aligned principal gradient per subject per session were calculated using the transformation matrices.

### 2.2. Aligned principal gradients take on an increasingly mixed character with increasing number of gradients used in Procrustes alignment

Next, we calculated correspondence between the aligned and unaligned principle gradient per subject per session as the ratio of the maximum to the sum of absolute values in the first column of the Procrustes transformation matrix (see Equation 2). This correspondence indicates the degree to which the first gradient after alignment corresponds to the one of the gradients before alignment. That is, if the correspondence value is close to 1, then one of the original gradients is essentially equivalent to the first gradient after alignment. The closer the correspondence value is to 0, the more the first gradient after alignment simply is a linear combination of all other gradients used in alignment. The correspondence between the aligned gradient and its unaligned version decreased with increasing number of gradients used for alignment (Fig. 1d). This indicates that the principal gradient takes on an increasingly mixed character by assimilating more information from lower gradients.

The decrease in correspondence reflects the increase in the fitting capacity of Procrustes alignment, i.e. accuracy of the alignment, which is a result of higher number of gradients used, essentially resulting in an “overfitted” aligned principal gradient. This is because the alignment can increasingly rely on information from the additional gradients rather than solely using the principal gradient to minimise the difference between the aligned subject-level gradient and the group-level (hold-out) template. These findings were highly consistent in the subsequent robustness analyses varying the parcellations (Schaefer 100, 200 and 400 parcellations), as well as kernel functions (spearman, pearson, normalised angle, gaussian, cosine, and no kernel) and dimensionality reduction approaches (diffusion map embedding, principal component analysis, laplacian eigenmaps) (see supplementary; Figs. S2 - S13). We also performed these analyses with the minimally processed HCP data to test if the effects could be exacerbated when more motion information is available. However the results remained reasonably similar (see supplementary; Fig. S14).

### 2.3. Procrustes Alignment correlates with Motion Signals

One possible source of subject-specific global information is motion in the scanner. We found the average framewise-displacement (FD) was correlated with the magnitude of the transformation (calculated as the sum of absolute values on the Procrustes transformation matrix; see Section 4.3). This correlation increased with an increasing number of gradients used for alignment (Fig. 2a). FD was also correlated with the magnitude of the transformation applied to specifically align the principal gradient (sum of absolute values of all elements in the first column of the transformation matrix; Fig. 2b). Importantly, this correlation was stronger and with a steeper increase when using the minimally processed fMRI data (for which motion regression and ICA-FIX were not applied) (Fig. 2a and b), indicating that Procrustes alignment has the capacity to use motion-related information if it is available.

**Figure 2:**
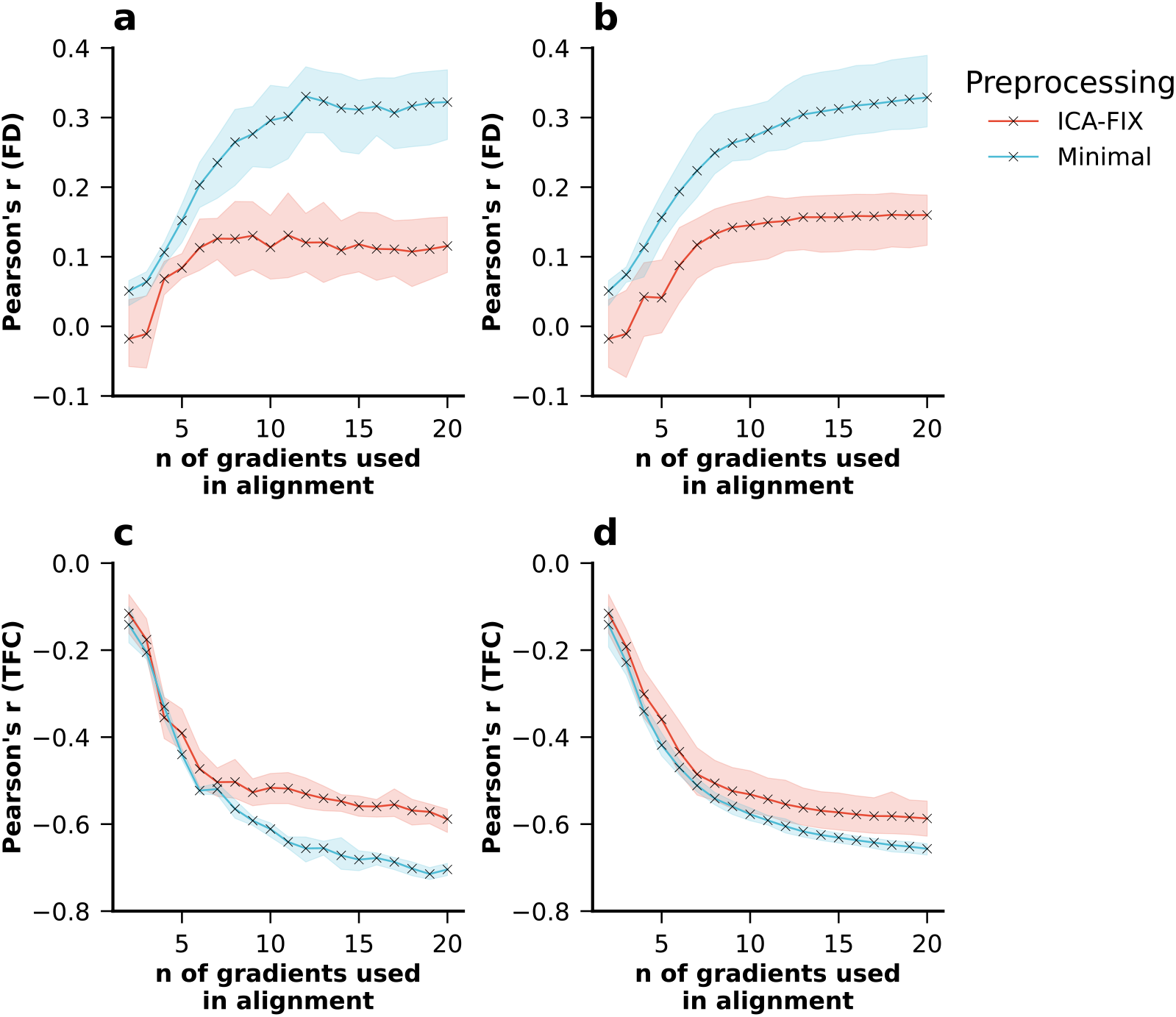
Mean Correlations across the 4 resting state sessions between in-scanner head motion and the magnitude of the transformation performed in Procrustes alignment for varying numbers of gradients used in alignment (fill areas indicate the standard deviation across the 4 resting state sessions). a) Correlation between FD and the sum of absolute values of all elements in the transformation matrix and b) correlation between FD and the sum of absolute values of all elements in the first column (i.e. the column that determines the alignment of principal gradient) of the transformation matrix. c) Correlation between TFC and the sum of absolute values of all elements in the transformation matrix and d) correlation between TFC and the sum of absolute values of all elements in the first column (i.e. the column that determines the aligned principal gradient) of the transformation matrix.

Additionally, typicality of functional connectivity (TFC; see Section 4.5.2) — an indicator of motion signal in the subjects’ FC compared to the group FC (Kopal et al., 2020) — was negatively correlated to the total transformation, and this relationship also became more pronounced with increasing number of gradients used in alignment (Fig. 2c). Similar effects are observed for the transformation magnitude specific to the principal gradient (Fig. 2d). Again, the relationship was stronger in fMRI data for which motion regression and ICA-FIX had not been applied. Overall, these findings suggest that the Procrustes transformation captures a higher degree of motion information when more gradients are used in alignment.

To test whether the motion signal present in the transformation matrix of the Procrustes alignment is due to inadequate preprocessing with leftover motion signal, we correlated each subject’s denoised, parcel-wise BOLD time series with the subject’s corresponding FD time series. For each session this yielded a normal distribution of correlations centered at 0 with a standard deviation of 0.03, indicating that the motion signal was appropriately removed from the BOLD time series (Fig. 3a-d). As expected, correlations between BOLD and FD time series were stronger for minimally processed data (see supplementary; Fig. S15)

**Figure 3:**
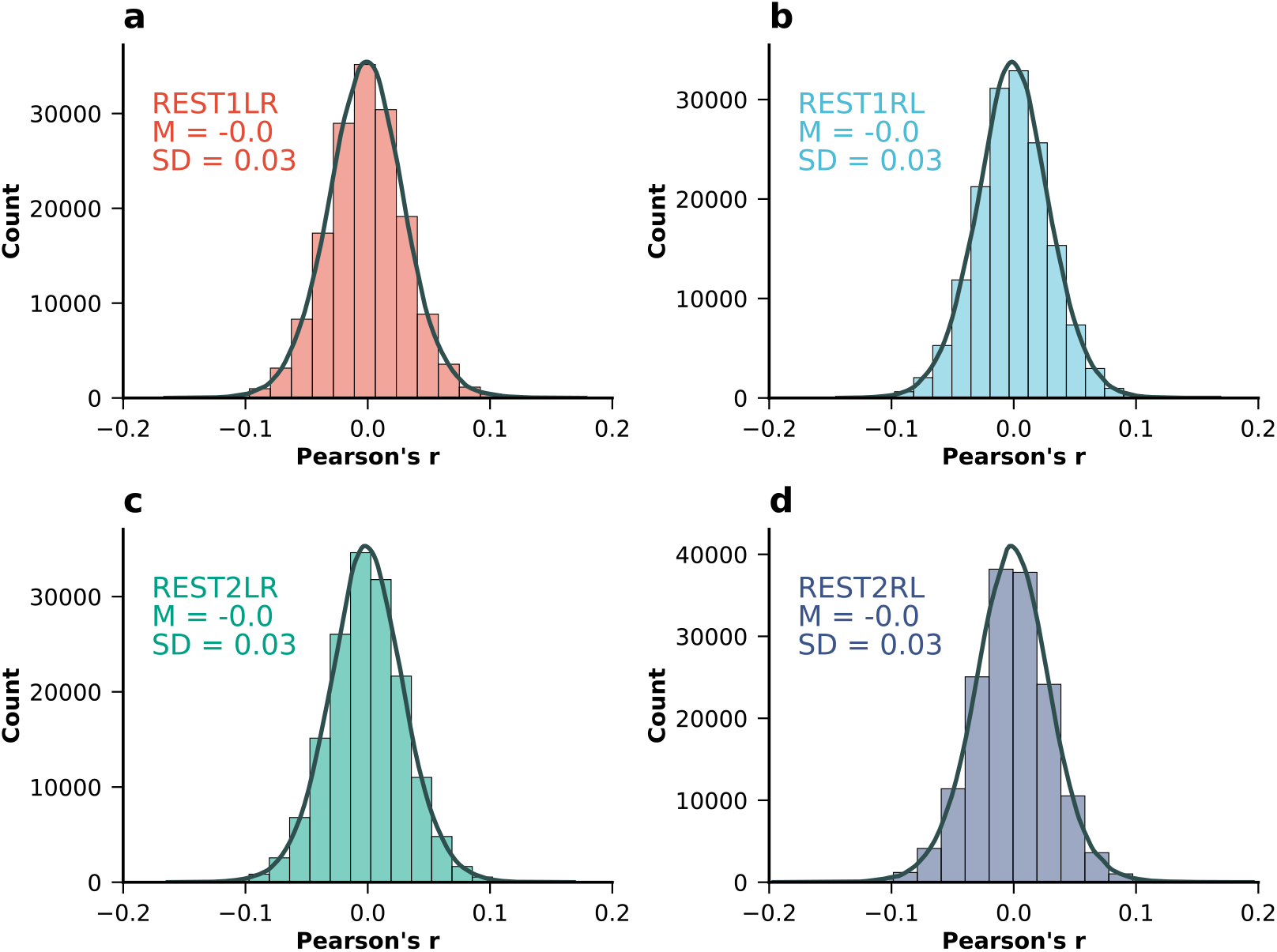
Distribution of correlations between each subjects’ ROI time series and FD time series for all four resting state fMRI sessions in the HCP-YA dataset: a) REST1LR, b) REST1RL, c) REST2LR, and d) REST2RL.

### 2.4. Procrustes alignment correlates with motion in the replication data sets

Next, we sought to replicate key findings regarding the correlation between in-scanner head motion and the magnitude of the transformation performed in Procrustes alignment, particularly in relation to the number of gradients used in alignment. In our analysis of the AOMIC PIOP1 and PIOP2 datasets, again, we found that correspondence between the unaligned and aligned principal gradient decreased with an increasing number of gradients used in alignment (Fig. 4a). However, we did not find the magnitude of the transformation to positively correlate with FD (Fig. 4b), but we found a strong negative correlation with TFC (Fig. 4c), indicating that the motion signal plays a role in Procrustes alignment of individual-level gradients. We further demonstrate that denoised subject’s ROI-wise BOLD time series do not correlate with motion in the PIOP1 (Fig. 4d) and in the PIOP2 (Fig. 4e) data sets.

**Figure 4:**
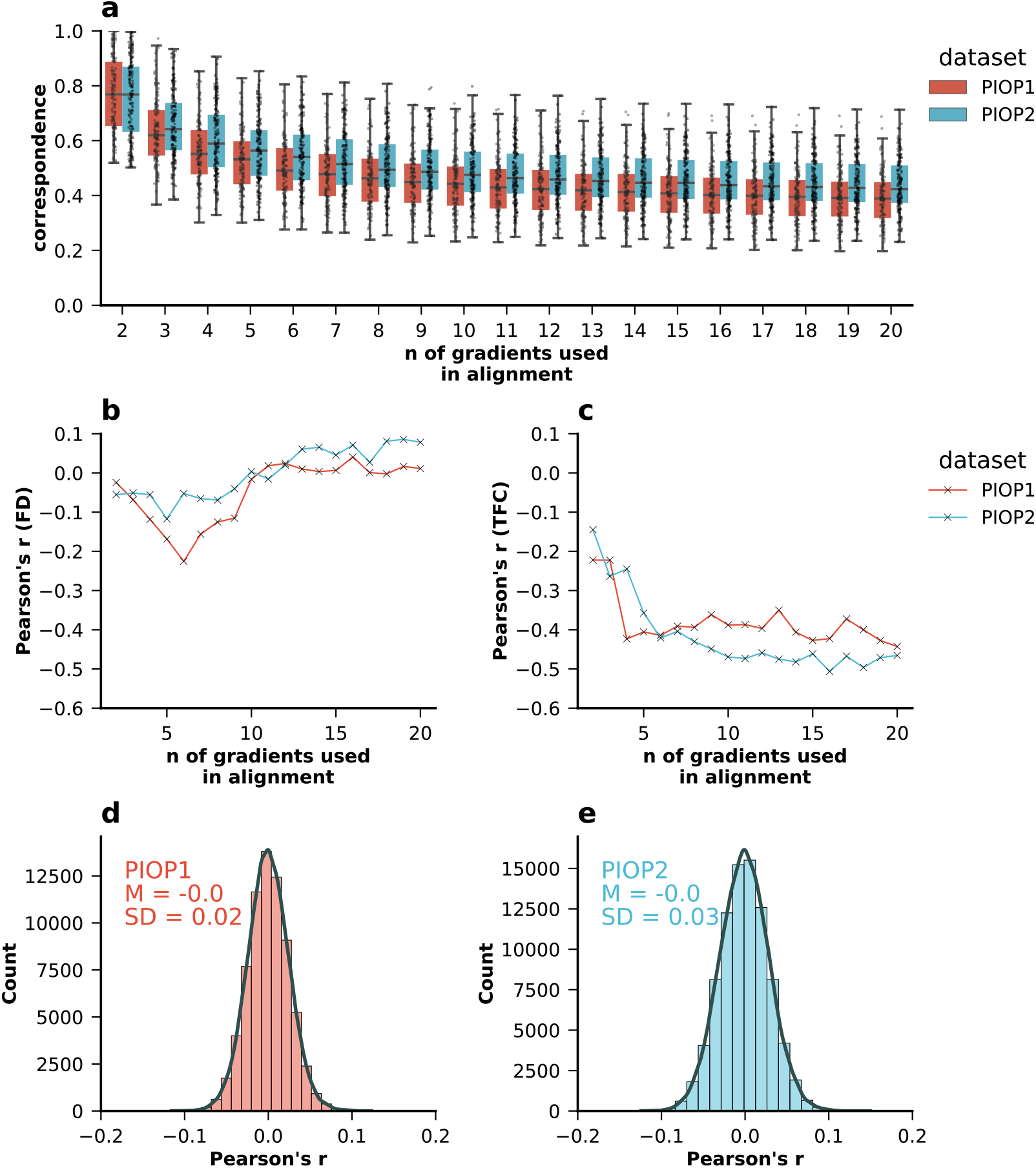
Replication of main analyses in the AOMIC PIOP1 and PIOP2 datasets: a) correspondence of the aligned principal gradient for varying number of gradients used in alignment, b) Pearson’s correlations between magnitude of the transformation (sum of all absolute values of the transformation matrix) and FD, c) Pearson’s correlations between magnitude of the transformation and TFC, d) Pearson’s correlations between denoised BOLD time series and FD time series in PIOP1 and e) PIOP2 datasets.

Since both HCP-YA as well as the AOMIC datasets have a rather narrow age distribution, and since age is related to head-motion in the scanner (Kato et al., 2020; Moqadam et al., 2024), we sought to replicate our findings in the Cam-CAN dataset, which has a wider age distribution (see Table 1). We found that correspondence again quickly decreased when increasing the number of gradients used in Procrustes alignment (Fig. 5a). We confirmed that the parcellated BOLD time series were not strongly related to FD (Fig. 5b). When extracting gradients and performing Procrustes alignment, the magnitude of the transformation was not strongly related to FD (Fig. 5c) but it was strongly related to TFC (Fig. 5d).

**Table 1:** Age and sex distribution for each of the 4 datasets used, and their respective “analysis” and “holdout” samples.

| Dataset | Sample | Count | Male | Female | M<sub>age</sub> | SD<sub>Age</sub> | MIN<sub>Age</sub> | MAX<sub>Age</sub> |
| --- | --- | --- | --- | --- | --- | --- | --- | --- |
| HCP-YA | **Whole** | **395** | **203** | **192** | **28.74** | **3.82** | **22** | **37** |
|  | Analysis | 296 | 152 | 144 | 28.83 | 3.71 | 22 | 37 |
|  | Holdout | 99 | 51 | 48 | 28.46 | 4.12 | 22 | 35 |
| AOMIC<br>PIOP1 | **Whole** | **175** | **78** | **97** | **22.09** | **1.77** | **18** | **26** |
|  | Analysis | 131 | 58 | 73 | 22.09 | 1.81 | 18 | 26 |
|  | Holdout | 44 | 20 | 24 | 22.02 | 1.66 | 18 | 26 |
| AOMIC<br>PIOP2 | **Whole** | **212** | **93** | **119** | **21.92** | **1.77** | **18** | **25** |
|  | Analysis | 158 | 70 | 88 | 21.81 | 1.76 | 18 | 25 |
|  | Holdout | 54 | 23 | 31 | 22.25 | 1.76 | 19 | 25 |
| Cam-CAN | **Whole** | **627** | **312** | **315** | **54.33** | **18.30** | **18** | **88** |
|  | Analysis | 470 | 234 | 236 | 53.8 | 18.72 | 18 | 88 |
|  | Holdout | 157 | 78 | 79 | 55.92 | 17.96 | 18 | 87 |

**Figure 5:**
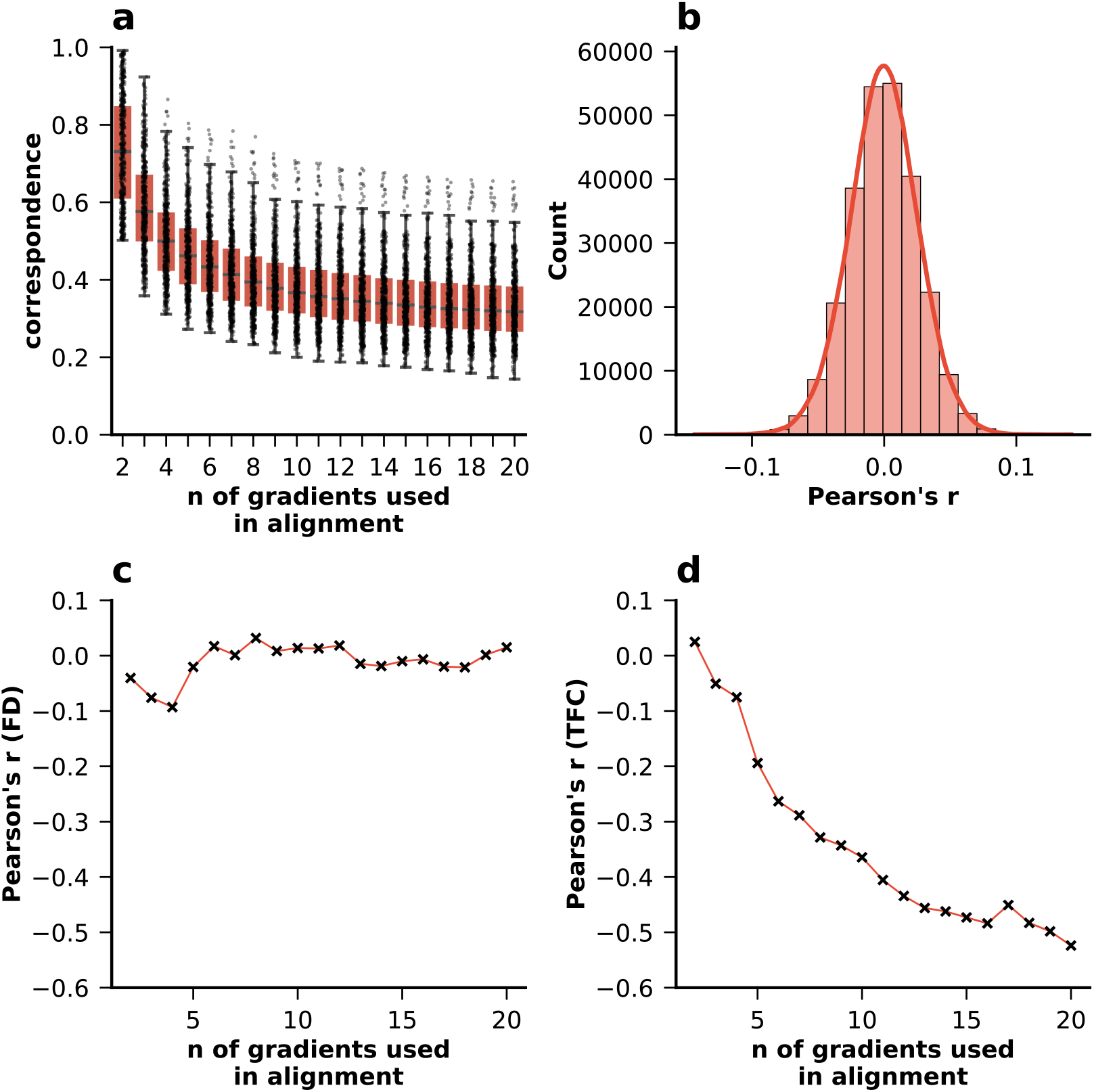
Replication of main analyses in the CamCAN dataset: a) correspondence of the aligned principal gradient for varying number of gradients used in alignment, b) Pearson’s correlations between denoised BOLD time series and FD time series, c) Pearson’s correlations between magnitude of the transformation (sum of all absolute values of the transformation matrix) and FD, d) Pearson’s correlations between magnitude of the transformation and TFC.

### 2.5. Procrustes alignment can impact the prediction of fluid intelligence and age

To examine the impact of Procrustes alignment on the prediction of commonly used targets in the FC literature, we sought to predict fluid intelligence, a psychometric measure which is available in all our datasets, using the aligned principal gradient. We could not find clear evidence of prediction in the HCP-YA (Fig. 6a), AOMIC PIOP1 (Fig. 6b), and PIOP2 (Fig. 6c) with *R*^2^ consistently close to 0. However, we found evidence that the principal gradient can predict fluid intelligence in the Cam-CAN dataset (Fig. 6d) with *R*^2^ (mean across folds and repeats) reaching a maximum value as high as 0.161 when using 14 gradients in Procrustes alignment. Further, prediction scores decreased drastically when removing FD as a confound, and when using FD and age together as confounds, *R*^2^ values were close to zero independent of the number of gradients used in alignment. In addition, we calculated the derivative of prediction scores to see the rate of change in prediction scores depending on the number of gradients used in alignment (Fig. 6e-h) and whether this rate of change was different between different confound removal scenarios. However, we did not find clear evidence that confound removal affected this rate of change.

**Figure 6:**
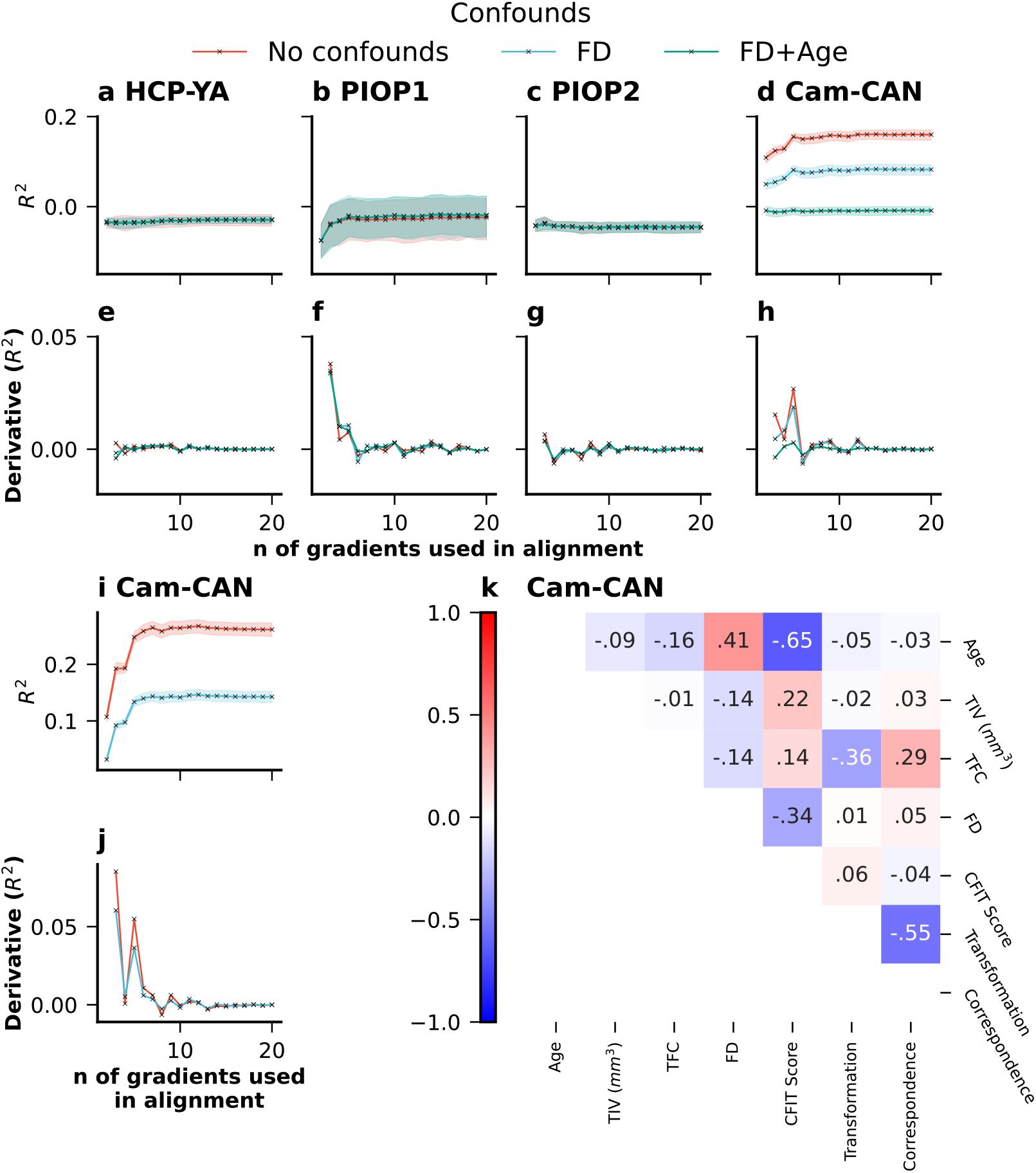
Prediction of fluid intelligence using the first principal gradient with varying numbers of gradients used in Procrustes alignment in a) HCP-YA (`PMAT24_A_CR`) b) AOMIC PIOP1 (`raven_score`), c) AOMIC PIOP2 (`raven_score`), and d) Cam-CAN (CFIT Score) as well as the corresponding derivatives of prediction scores (*R*^2^) in e) HCP-YA, f) AOMIC PIOP1, g) AOMIC PIOP2, and d) Cam-CAN datasets. i) Prediction of age in the Cam-CAN dataset as well as the j) corresponding derivative of prediction scores (*R*^2^). k) Correlations between the transformation magnitude, correspondence (when using 10 gradients in Procrustes alignment), CFIT Score, FD, TFC, TIV, and Age in the Cam-CAN dataset.

Since this result clearly indicated a relationship between the principal gradient and age, and because age is a common prediction target in neuroimaging research in its own right, we also sought to use the principal gradient to predict age in the Cam-CAN dataset. We found again that increasing the number of gradients used in Procrustes alignment led to higher prediction scores (Fig. 6i) with a mean R2 as high as 0.268 when using 12 gradients in Procrustes alignment. Again, we looked at the rate of change, and this time, we did observe that not removing FD as a confound initially led to slightly higher increases of the prediction scores (Fig. 6j). We examined the Cam-CAN dataset, finding strong negative correlations between age and fluid intelligence, a positive correlation between age and motion, and a weak negative correlation between age and TFC, with TFC strongly linked to Procrustes alignment transformation magnitude using 10 gradients (Fig. 6k). In addition, we compared the distributions of average FD values between the four datasets (and the individual HCP-YA sessions), and as expected (due to the wider age range), we find that FD values are overall larger and more dispersed in the Cam-CAN dataset (see supplementary; Fig. S16).

### 2.6. Prediction of motion signal

To assess the degree to which the number of gradients in Procrustes alignment affects the motion signal present in the principal gradient, we also used a machine learning approach to classify subjects as “high-” or “low-motion” subjects using aligned principal gradients as features. We binarised FD using median FD as a threshold, as we expected a weak signal, and classification tasks are typically easier to solve than regression (Greene et al., 2022). Across all four sessions in the HCP-YA dataset, classification accuracy and the area under the receiver operating characteristic curve (AUC) were close to the chance-level, indicating little to no evidence of successful out-of-sample prediction (Fig. 7a and b).

**Figure 7:**
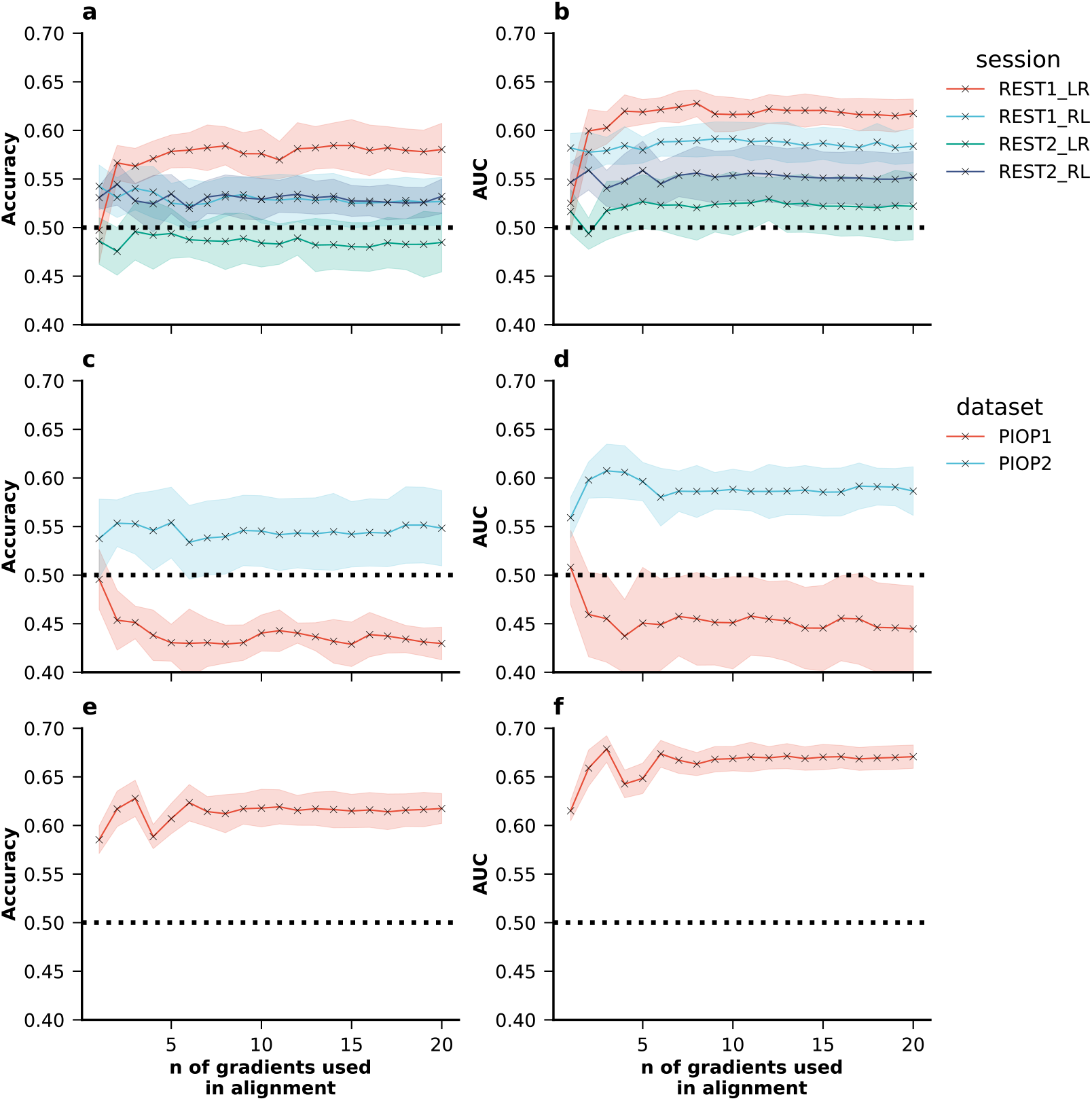
a) Classification accuracy and b) area under the receiver operating characteristic curve for classification of “high-” and “low-motion” subjects in the HCP-YA, c) classification accuracy and d) area under the receiver operating characteristic curve in the AOMIC PIOP1 and AOMIC PIOP2, and e) classification accuracy and f) area under the receiver operating characteristic curve in the Cam-CAN datasets using ridge classifiers. The dashed black line indicates chance level.

Similar chance-level performance was found in both the AOMIC PIOP1 dataset and the AOMIC PIOP2 dataset (Fig. 7c and d). In the Cam-CAN dataset, with the widest age distribution out of these datasets, we found evidence that classification of high-vs low-motion subjects was above the chance-level (Fig. 7e and f). This result may be attributed to the diverse age distribution in the Cam-CAN dataset, which could introduce additional variability that aids in the classification process. However, in none of the datasets we found a clear pattern of classification accuracy increase or decrease depending on the number of gradients used in Procrustes alignment.

## 3. Discussion

In this study, we investigated the impact of Procrustes alignment on the aligned principal gradient and its implications for subject-level downstream analyses. Specifically, we evaluated the impact of the number of gradients used in alignment. Our findings shed light on several critical aspects regarding the application and interpretation of Procrustes alignment in individual-level functional gradient studies. The results demonstrate that the aligned principal gradient incorporates information not just from the unaligned principal gradient but from all other gradients. We observed a decreasing correspondence between aligned and unaligned principal gradients with an increasing number of gradients, confirming the idea that the aligned principal gradient becomes increasingly influenced by other gradients, taking on a more mixed character. A similar effect has been reported in the application of Procrustes rotation to PLS permutation testing, and as pointed out by others, it will likely extend to further applications of Procrustes rotation as well (Danyluik et al., 2024).

Our results suggest that motion signals are introduced to the aligned principal gradient through the Procrustes alignment. We observed a positive correlation between average FD and the magnitude of Procrustes transformation, which intensified with an increasing number of gradients used for alignment Fig. 2. This finding is further strengthened by a FC-derived motion measure (TFC) which showed a negative correlation with the magnitude of transformation indicating that lower TFC, associated with higher motion, corresponds to a greater degree of transformation in Procrustes alignment. The correlation between the transformation and TFC was higher than that with FD and was more consistent across datasets. This is expected considering that the purpose of the Procrustes alignment is to make subject-gradients more typical. Overall, our results suggest that the Procrustes alignment incorporates a higher degree of motion information as more gradients are used for Procrustes alignment.

The number of gradients used in Procrustes alignment strongly influenced identification accuracy and differential identifiability of the principal gradient. As the number of gradients increased, identification accuracy improved while differential identifiability decreased. However, as we also point out, differential identifiability behaves more consistently with identification accuracy after applying Fisher’s r-to-z transform to correlation values. This finding may explain why previous research has occasionally found some inconsistencies in the behaviour of identification accuracy and differential identifiability (Cutts et al., 2023; Sasse et al., 2023). Overall, the impact of the number of gradients used in alignment on identification and identifiability can be explained by the observation that the aligned principal gradient integrates information from all gradients used in the alignment process. This is in particular demonstrated by the decrease of correspondence — the degree to which the first gradient after alignment corresponds to one of the gradients before alignment — when increasing the number of gradients used in alignment.

Our machine learning analysis corroborates these findings to some degree. While we did not find evidence of prediction of fluid intelligence using the principal gradient in the HCP-YA and AOMIC datasets, fluid intelligence could be predicted in the Cam-CAN dataset with *R*^2^ consistently above 0.1. This level of accuracy is plausible given that a FC-based meta-analysis has found Pearson’s r between true and predicted fluid intelligence close to 0.15 (Vieira et al., 2022). Studies using the Cam-CAN dataset specifically have found prediction scores even above that, likely also due to confounding effects of age, but also possibly due to its use of Cattels’ Culture Fair Intelligence Test (CFIT) rather than Raven’s Progressive Matrices (PMAT) as used in the HCP and AOMIC datasets (Jiang et al., 2022). More importantly, we found that increasing the number of gradients used in Procrustes alignment increased prediction of fluid intelligence in the Cam-CAN dataset, but that removal of age and motion information (average FD) also removed the principal gradients capacity to predict fluid intelligence independent of the number of gradients used in alignment. Similarly, we found that we could successfully predict age using the principal gradient in the Cam-CAN dataset, and that increasing the number of gradients used in Procrustes alignment also increased prediction scores. Importantly, when removing the confounding effect of average FD, prediction scores dropped and in addition the increase in prediction scores with each additional gradient used in alignment decreased slightly. This indicates that in-scanner head motion contributes to better prediction success when more gradients are used in alignment.

Lastly, we attempted to predict the subject-level head motion using the prinicpal gradient. We expected better prediction success with more gradients used in Procrustes alignment. While we did not find clear evidence of successful classification of motion in the HCP-YA and AOMIC datasets, we found good evidence of successful classification in the Cam-CAN dataset. This may be a result of the relatively low sample sizes in the former datasets which is not favourable for machine learning analysis. In contrast, the Cam-CAN dataset has the largest sample size as well as more variance of age and FD. This is important since age is related to head-motion in the scanner (Kato et al., 2020; Moqadam et al., 2024) as well as gradients (Bethlehem et al., 2020) and a wider age range is therefore likely to improve gradient-based prediction of motion. However, even in the Cam-CAN dataset we did not find evidence for increased classification accuracy with more gradients used in Procrustes alignment.

One limitation affecting our ability to predict motion in the classification analysis is the inherent dilution of the motion signal accumulated over multiple stages of analysis. FD, commonly used as a measure of motion, is derived indirectly from the fMRI time series, where each time point reflects a snapshot of head position changes across translational and rotational dimensions. As a result, FD captures movement indirectly rather than directly reflecting physiological or anatomical features related to motion. Furthermore, by averaging FD values across all time points for each subject, the dynamic variability of motion over the scanning session is reduced to a single summary statistic, which can mask significant temporal fluctuations and lead to further signal loss. These factors mean that the motion signal is both indirect and reduced, limiting the ability of our classification models to accurately classify subjects based on their average FD.

In addition, although FD provides a useful proxy for in-scanner head motion, it captures only a partial aspect of motion-related artefacts (Goto et al., 2015). Importantly, FD is a catch-all measure, and similar FD values can have multiple underlying types of motion (Power et al., 2014). Moving forward, future studies could benefit from incorporating more direct measures of motion using external cameras or motion-tracking devices (Kim et al., 2013; Seto et al., 2001). These technologies can provide real-time, high-resolution data on head movements, allowing for a more comprehensive understanding of motion artefacts during MRI scans.

These results have significant implications for the interpretation and application of Procrustes alignment in functional gradient studies, especially for individual-level analysis. The observed relationship between alignment characteristics and motion signals underscores the importance of considering motion artefacts in FC analyses, particularly in individual-level studies where motion can significantly impact data quality and interpretation. For example, it has been found that motion signals reliably correlate with behavioural outcomes (Bolton et al., 2020; Siegel et al., 2017). Additionally, some strategies to account for motion (e.g. excluding high-motion subjects) can bias datasets and lead to understudying of important clinical populations (Cosgrove et al., 2022; Nebel et al., 2022). Therefore, rather than excluding subjects with high head-motion, it is often suggested to employ appropriate strategies to address motion artefacts, ensuring that analyses remain robust and inclusive (Kay et al., 2023; Parkes et al., 2018; Satterthwaite et al., 2019). To this end, it may also be worthwhile exploring the prospect of alternative gradient alignment solutions.

Moreover, the findings emphasise the need for careful consideration of the number of gradients used in alignment, as it directly influences the characteristics and interpretability of the aligned principal gradient. While using a higher number of gradients can increase the fit between group- and individual-level gradients (Hardikar et al., 2024; Mckeown et al., 2020), this comes as a trade-off in terms of validity: the benefit in terms of better fit may stem from undesired signals captured by lower gradients. Intuitively, aligning with a high number of gradients may be thought of as a kind of “overfitting”. In our study we further demonstrate that motion signals are likely introduced into the principal gradient via Procrustes alignment performed with higher numbers of gradients. As a result it may not be desirable to recommend a specific number of gradients to be used in alignment, but to consider the advantages of a better group-level fit and the disadvantages it may incur in terms of the introduction of undesired signals together with the specific characteristics of the dataset.

To some degree our results showcase one of the reasons for extracting gradients in the first place: identify gradients that presumably capture neural signals, and discard those that capture other sources of information and noise (Bijsterbosch et al., 2021). Simply aligning a specific number of gradients risks introducing undesirable signals back into the gradients of interest. It may be more beneficial to make the choice of number of gradients to align based on specific biological models (Bolt et al., 2022; Margulies et al., 2016; Mesulam, 1998; Zhang et al., 2019) rather than using an ad-hoc predefined number (Hardikar et al., 2024; Mckeown et al., 2020). Researchers can also perform robustness analyses that test whether their results hold for different numbers of gradients used in Procrustes alignment. These robustness checks can be informative whether the impact of alignment on the results is practically meaningful. In addition, of course, researchers should report the exact number of gradients used in Procrustes alignment, and ideally report whether they did or did not try their analysis with varying numbers of gradients. Additionally, it is important to consider head motion as a confound in studies of functional gradients, and determining the most effective approach to address this remains an active area of research (Kay et al., 2023).

In conclusion, our study provides valuable insights into the impact of Procrustes alignment on individual-level functional gradient analyses, highlighting its role in integrating information from multiple gradients and its relationship with motion signals. Future research can investigate the mechanisms underlying Procrustes alignment and explore strategies to mitigate the effects of motion artefacts in FC gradient studies, ultimately enhancing our understanding of the human brain’s functional organisation.

## 4. Methods

### 4.1. Datasets

For this project, we used 4 different datasets: the Human Connectome Project – Young Adults (HCP-YA) dataset, the Amsterdam Open MRI Collection (AOMIC) Population Imaging of Psychology (PIOP) datasets 1 and 2, and lastly the Cambridge Centre for Ageing and Neuroscience (Cam-CAN) dataset. Each dataset was split into two partitions: A holdout sample consisting of 20% of the data was used to create a group average gradient that could be used as a reference for Procrustes alignment, whereas 80% of the data was used to create individual-level gradients for subject-level analyses. Age, sex, as well as the partitioning of the data can be seen in Table 1. Retrospective analysis of these datasets was approved by the local Ethics Committee at the Faculty of Medicine at Heinrich-Heine-University in Düsseldorf (2018-317-RetroDEuA).

#### 4.1.1. Human Connectome Project - Young Adult

For the primary, exploratory analyses in this project, we used the HCP-YA dataset. The details regarding collection of behavioural data, fMRI acquisition, and image preprocessing in the HCP-YA have been described elsewhere (Barch et al., 2013; Glasser et al., 2013; Van Essen et al., 2013). Here we aim to give a brief overview. The scanning protocol for the HCP-YA was approved by the local Institutional Review Board at Washington University in St. Louis. We used data obtained from four resting-state fMRI (rs-fMRI) sessions.

The four sessions of rs-fMRI were obtained on two separate days (each lasted ca. 15 minutes; 60 minutes across all four sessions). On each day, two sessions were recorded for different phase encoding directions (left-right [LR] and right-left [RL]) providing four overall rs-fMRI datasets. Scans were acquired using a 3T Siemens connectome-Skyra scanner with a gradient-echo EPI sequence (TE=33.1ms, TR=720ms, flip angle = 52°, 2.0mm isotropic voxels, 72 slices, multiband factor of 8). This data had already undergone the HCP’s ICA-FIX (independent component analysis and FMRIB’s ICA-based X-noiseifier) procedure (Salimi-Khorshidi et al., 2014), which also included removal of Friston 24 motion parameters (Glasser et al., 2013). In addition, we regressed out white matter (WM), cerebro-spinal fluid (CSF), and global signals (GS), their squared terms, and temporal derivatives. The data was bandpass filtered at 0.01 - 0.08 Hz.

#### 4.1.2. Amsterdam Open MRI Collection

We used two datasets from the Amsterdam Open MRI Collection (AOMIC), known as the Population Imaging of Psychology (PIOP) data sets 1 and 2 (Snoek et al., 2021) to replicate and validate our main findings. The exact data collection protocols are described elsewhere in detail, so here we provide a short overview. The University of Amsterdam’s ethical committee approved these studies before data collection started (PIOP1 EC number: 2015-EXT-4366, PIOP2 EC number: 2017-EXT-7568).

The functional MRI scanning protocol for the PIOP1 and PIOP2 data sets utilised a Philips 3 T scanner, with PIOP1 scanned on the “Achieva’’ version and PIOP2 on the “Achieva dStream” version, both equipped with a 32-channel head coil. Functional MRI data in PIOP1 for resting-state fMRI were acquired with multiband acceleration, whereas functional MRI scans in the PIOP2 were acquired sequentially. For both data sets, resting state scans lasted 6 minutes (PIOP1; i.e., 480 volumes with a 0.75 second TR) and 8 minutes (PIOP2; i.e., 240 volumes with a 2 second TR).

We used data processed using fMRIPrep as provided by the original PIOP studies (Esteban et al., 2019; Snoek et al., 2021). Spatial normalisation to the ICBM 152 Nonlinear Asymmetrical template was performed using nonlinear registration with ANTs. For functional data, motion correction was conducted using mcflirt, and “fieldmap-less” distortion correction was applied by co-registering functional images to T1-weighted images with intensity inversion. Similar to the HCP-YA dataset, we regressed out the Friston 24 motion parameters, white matter (WM), cerebro-spinal fluid (CSF), and global signals (GS), their squared terms, and temporal derivatives. The data was bandpass filtered at 0.01 - 0.08 Hz.

#### 4.1.3. Cambridge Centre for Ageing and Neuroscience

Because both HCP-YA and the AOMIC datasets have a rather narrow age distribution with mainly young adults, we also sought to replicate our findings using resting state fMRI data from the Cam-CAN dataset. Collection of this data has been approved by the Cambridgeshire 2 Research Ethics Committee (reference: 10/H0308/50). Data from Cambridge Centre for Ageing Neuroscience dataset were retrieved from their official source at the CamCAN Data Portal (https://camcan-archive.mrc-cbu.cam.ac.uk/dataaccess/) (Shafto et al., 2014; Taylor et al., 2017) and provided in the form of a DataLad dataset, a research data management solution providing data versioning, data transport, and provenance capture (Halchenko et al., 2021).

Resting state fMRI data in the Cam-CAN dataset were acquired using a T2*-weighted Gradient-Echo Echo-Planar Imaging (EPI) sequence while participants rested with their eyes closed. A total of 261 volumes were collected, each comprising 32 axial slices acquired in descending order. The slices had a thickness of 3.7 mm with an interslice gap of 20%, ensuring whole-brain coverage including the cerebellum. The acquisition parameters included a repetition time (TR) of 1970 milliseconds, an echo time (TE) of 30 milliseconds, a flip angle of 78 degrees, and a field of view (FOV) of 192 mm × 192 mm. The voxel size was 3 mm × 3 mm × 4.44 mm, and the total acquisition time for each scan was 8 minutes and 40 seconds.

In order to keep preprocessing as consistent as possible, we used fMRIPrep with similar configurations to the AOMIC datasets. Spatial normalisation to the ICBM 152 Nonlinear Asymmetrical template was performed using nonlinear registration with ANTs. For functional data, motion correction was conducted using mcflirt. Again, we regressed out the Friston 24 motion parameters, white matter (WM), cerebro-spinal fluid (CSF), and global signals (GS), their squared terms, and temporal derivatives. The data was bandpass filtered at 0.01 - 0.08 Hz.

### 4.2. Gradient Extraction and Alignment

Denoised time series were aggregated using the Schaefer 400 parcellation (Schaefer et al., 2018), a popular choice for FC-based prediction and functional gradient studies (Hardikar et al., 2024; He et al., 2020; Kong et al., 2019). FC was calculated as pairwise Pearson’s correlation between ROI time series resulting in a 400×400 connectivity matrix per subject per session. For subjects in the main analysis dataset, gradients for each session were extracted from these FC matrices using the BrainSpace toolbox (Wael et al., 2020), which also provides Procrustes alignment to find the optimal transformation matrix to minimise the sum of squared errors of a source matrix to a reference matrix. Here, we focus on gradient extraction using the following recommended parameters: kernel = normalized_angle, sparsity = 0.9, dimensionality reduction technique = diffusion map embedding (“dm”). These parameters are among the most commonly used in the literature (Bethlehem et al., 2017; 2020; Hänisch et al., 2023; Kong et al., 2023; Margulies et al., 2016; Wan et al., 2023). In addition, as a robustness check, we provide results for alternative parameters in the supplement.

For subjects in the 20% holdout dataset, FC matrices were averaged after applying Fisher’s r-to-z transform. Elements in this average matrix underwent the reverse transform to derive correlation values and using this average FC matrix an average reference gradient set was derived for alignment. The gradients of subjects in the analysis dataset were then each aligned to this holdout reference gradient using Procrustes alignment. Subsequent downstream analyses were performed using these aligned gradients. The reference was derived separately for each of the four datasets.

### 4.3. Procrustes Alignment and Correspondence of Aligned and Unaligned Principal Gradients

Before comparing gradients, it is important to align them for two main reasons: Firstly, the subject-level and group-level gradients can be in a different order, e.g. the second subject-level gradient might match better with the reference principal gradient. Secondly, the sign of parcels within gradients can also vary arbitrarily across subjects. Procrustes alignment aims to solve this by linearly transforming a given set of unaligned gradients to optimally align with a set of reference gradients. For example, let *A* be a *m* × *n* matrix containing a subject’s *n* gradients consisting of *m* ROIs, and *B* be the equivalent *m* × *n* reference gradient matrix. Procrustes alignment is then performed using an *n* × *n* transformation matrix *T* to transform *A* such that the sum of squared errors between *B* and the transformed matrix *A*^′^ is minimised, i.e. *A*^′^ = *AT*.

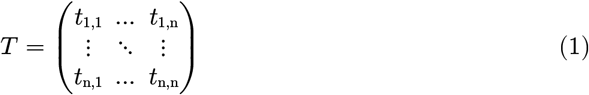

Since the aligned principal gradient is a linear combination of the original, unaligned gradients, the transformation matrix provides information about each unaligned gradients’ contribution toward the aligned principal gradients. To quantify this, we calculated the relative magnitude of the unaligned principal gradient with respect to all gradients used for alignment. If the contribution of the unaligned principal gradient toward the aligned principal gradient is relatively large, then we can conclude that the unaligned and the aligned principal gradient have a high correspondence with each other.

This “Correspondence” is calculated as the ratio of the maximum to the sum of absolute values in the first column of the transformation matrix *T* which contains the weights for each original unaligned gradient toward the aligned principal gradient. That is, *t*_1,1_ represents the contribution of the first unaligned gradient toward the aligned principal gradient, and more generally, *t*_i,1_ represents the contribution of the *i*-th unaligned gradient toward the aligned principal gradient. More formally, to quantify the correspondence between the unaligned and aligned principal gradients (i.e. the first column of the untransformed original matrix *A* and the transformed matrix *A*′), we calculate the correspondence as follows:

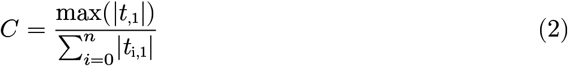

Here |*t*_i,1|_ represents the absolute value of each element in the first column. This expression accounts for possible reordering of the unaligned gradients as well as change in the sign. In other words, *C* quantifies how dominant the largest weight is compared to the overall weights in the first column of the transformation matrix *T*. It quantifies how dominantly the unaligned principal gradient determines the aligned principal gradient compared to all other gradients. That is, if *C* is close to 1, it suggests that the transformation is dominated by the unaligned principal gradient indicating its strong correspondence with the aligned principal gradient. Conversely, if *C* is closer to 0, it implies a more distributed adjustment using all unaligned gradients, indicating a weaker correspondence.

We also calculated the magnitude of the Procrustes rotation as the sum of absolute values in the transformation matrix, to get a sense of the degree to which the rotation conveys information on subject motion. This sum provides a quantitative measure of the extent of alignment required to match the subject data to the reference. To further dissect the impact of Procrustes rotation, we specifically focused on the alignment of the principal gradient—a key component in functional gradient analysis. For this, we calculated the magnitude of the transformation applied to align the principal gradient as the sum of absolute values in the first column of the transformation matrix. This column determines the transformation of the principal gradient during Procrustes alignment, and the magnitude of its summed elements can thus indicate how much the principal gradient is altered to achieve alignment. A higher sum would suggest a greater transformation, implying that the subject’s brain data diverged more from the reference, possibly due to subject motion or intrinsic variability.

### 4.4. Identification and Differential Identifiability

We performed identification (Finn et al., 2015) and calculated differential identifiability (Amico & Goñi, 2018) to test the impact of varying the number of gradients used in Procrustes alignment on downstream subject-level analyses. Identification was performed by utilising an individual’s principal functional gradient derived from a resting state fMRI scanning session to find the best match from a database of principal functional gradients derived from another resting state fMRI scanning session. The identification accuracy is determined by the proportion of participants correctly identified. Differential identifiability is determined in a similar fashion by correlating each subject’s principal functional gradient derived in one fMRI scanning session, with each subject’s principal gradient derived in another fMRI scanning session. Differential Identifiability between those two sessions is then defined as the difference between mean within-subject correlations (*I*_self_) and mean between-subject correlations (*I*_other_): *I*_Diff_ = (*I*_self_ − *I*_other_) × 100. However, simply subtracting *I*_other_ from *I*_self_ may be problematic as Pearson’s r is not a distance metric, because it does not measure distance in a way that would allow for straightforward comparison or summation. For instance, Pearson’s r is not strictly equidistant, meaning that the “distance” between two points in the metric space does not consistently reflect their relative closeness. In other words, it may fail to maintain equal spacing between pairs of points that intuitively would be considered equally distant (Solo, 2019). Therefore we performed an additional analysis in which Fisher’s r-to-z transform is first applied to all correlation values before computing the difference. Identification accuracy and differential identifiability were calculated for each pairwise combination of sessions in the HCP-YA dataset (N_Sessions_ = 4; N_Combinations_ = 6).

### 4.5. Motion Signal

#### 4.5.1 Framewise Displacement

Motion was characterised using average framewise displacement (FD) for each subject. FD is a composite measure derived from six head motion parameters, encompassing translational displacements along the X, Y, and Z axes, as well as rotational displacements of pitch, yaw, and roll, transitioning from one volume to the subsequent (Power et al., 2012). It represents the mean disparity among rotation and translation parameters. FD was calculated for each volume in the fMRI data and then averaged to obtain an aggregate measure of motion for the session.

#### 4.5.2. Typicality of Functional Connectivity

In addition to examining motion through FD, we incorporated the concept of typicality of functional connectivity (TFC) as an additional measure of motion derived directly from FC which has been shown to negatively correlate with measures of head motion in the scanner. While FD captures motion within a subject based on differences between volumes, TFC offers a complementary perspective by assessing motion across subjects relative to the group mean (Kopal et al., 2020). In addition, FD is derived from the raw time series, whereas TFC serves as a marker of motion of the FC directly, after all processing and denoising has taken place. TFC is defined as:

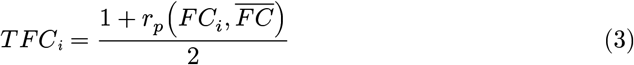

where *r*_*p*_ is Pearson’s correlation, *FC*_*i*_ a subject’s FC, and 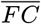 is the group average (i.e. typical FC). In this study, this group average FC was derived from the 20% holdout (see Table 1).

### 4.6. Prediction of Fluid Intelligence and Age

To investigate the impact of Procrustes alignment on common machine learning workflows, we attempted to predict fluid intelligence across all datasets, and age specifically in the Cam-CAN dataset due to its diverse age range (see Table 1). As a primary model, we used a ridge regression as implemented by scikit-learn (Pedregosa et al., 2011). To estimate the generalisation error of the predictive model, we employed 10 times repeated 5-Fold nested cross-validation (CV). Within each nested CV, an inner 5-Fold cross-validation was used on each train partition of the respective splits to determine the optimal hyperparameters. We tuned the `alpha` parameter using a grid generated by numpy (Harris et al., 2020) as `np.geomspace(0.001, 10000, 50)`. In the HCP-YA and AOMIC datasets, we used Raven’s Progressive Matrices (`PMAT24_A_CR` in the HCP-YA and `raven_score` in the AOMIC datasets). In the Cam-CAN dataset we used Cattel’s Culture Fair Intelligence Test (CFIT) Total Score. In the prediction of fluid intelligence we followed three confound removal scenarios: 1) Remove no confounds, 2) remove FD as a confound, and 3) remove FD and age as a confound. In the prediction of age we followed two confound removal scenarios: 1) Remove no confounds, and 2) remove FD as a confound. Each time confounds were removed in a cross-validation consistent way, i.e. by training the confound removal model on the train partition only, and then applying it to the test partition as To investigate the individual-level motion-related information present in the principal gradient after Procrustes alignment, we used machine learning to predict the subjects’ head-motion using the aligned principal gradient as features. To this end we binarized FD scores to create unambiguous classification outcomes using median FD as threshold, such that subjects with below median FD were considered “low-motion” subjects, whereas subjects with above median FD were considered “high-motion” subjects. A similar binarization approach has been proposed previously (Greene et al., 2022). As a primary model, we used a ridge classifier as implemented by scikit-learn (Pedregosa et al., 2011). In addition, we performed this analysis using a linear support vector machine (SVM) as well as SVMs with a radial basis function (RBF) kernel as well as random forest (RF) classifiers to reproduce the results using non-linear models.

To estimate the generalisation error of the predictive models, we employed 10 times repeated 5-Fold nested cross-validation (CV). Within each nested CV, an inner 5-Fold cross-validation was used on each train partition of the respective splits to determine the optimal hyperparameters. For the ridge classifier we tuned the `alpha` parameter using a grid generated by numpy (Harris et al., 2020) as `np.geomspace(0.001, 10000, 50)`. For the linear SVM, we generated a grid to optimise the `C` parameter (`np.geomspace(1e-4, 1e4, 10)`). In the RBF SVM we additionally optimised the parameter of the RBF kernel (`np.geomspace(1e-9, 1e4, 10)`). For the RF classifier, we tuned the `max_depth` parameter using the grid `[5, 10, 20, None]`. After hyperparameter selection a model was fitted on the whole train partition and then tested on the left-out test samples of the CV split.

### 4.7. Software

All code written for this project is made publicly available here: https://github.com/juaml/procrustes_alignment_for_fc_gradients/. The code for this project was written and executed using Python (version 3.11.4) and its scientific ecosystem, in particular using the following software packages: numpy (version 1.25.2; (Harris et al., 2020)), pandas (version 1.5.3; (McKinney, 2010)), scikit-learn (version 1.2.2; (Pedregosa et al., 2011)), scipy (version 1.11.2; (Virtanen et al., 2020)), julearn (version 0.3.0; (Hamdan et al., 2024)), junifer (version 0.0.3; (Mandal et al., 2023)), matplotlib (version 3.7.2; (Hunter, 2007)), and seaborn (version 0.12.2; (Waskom, 2021)). In addition, we forked the latest release version (0. 1.10) of BrainSpace (Wael et al., 2020) in order to adjust the code to make transformation matrices generated in Procrustes alignment available to the user. This fork is available at https://github.com/LeSasse/BrainSpace_release.

## 4.8. Acknowledgments

This research was partially funded by the Helmholtz Portfolio Theme “Supercomputing and Modelling for the Human” and by the Max Planck School of Cognition supported by the Federal Ministry of Education and Research (BMBF) and the Max Planck Society (MPG). Data collection and sharing for this project was provided by the Cambridge Centre for Ageing and Neuroscience (CamCAN). CamCAN funding was provided by the UK Biotechnology and Biological Sciences Research Council (grant number BB/H008217/1), together with support from the UK Medical Research Council and University of Cambridge, UK.

## Notes

**Funding** This research was partially funded by the Helmholtz Portfolio Theme “Supercomputing and Modelling for the Human” and by the Max Planck School of Cognition supported by the Federal Ministry of Education and Research (BMBF) and the Max Planck Society (MPG).

https://github.com/juaml/procrustes_alignment_for_fc_gradients

### Competing Interest Statement

The authors have declared no competing interest.

