## Supplementary Figures for "Procrustes Alignment in Individual-level Analyses of Functional Gradients"

November 26, 2024

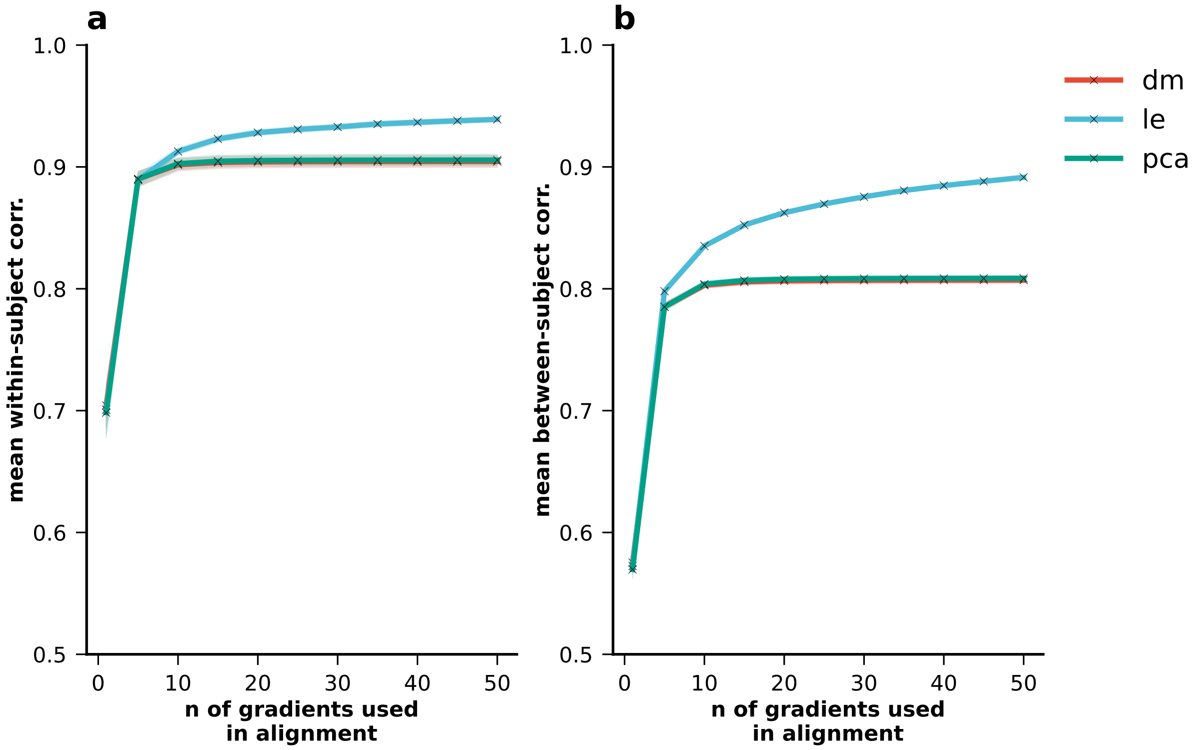


**Figure S1.** Impact of Procrustes alignment on a) mean within-subject correlations and b) mean between-subject correlations. For each subject, gradients were extracted per session (kernel = normalized_angle; sparsity = 0.9). They were then aligned to the holdout reference gradient using Procrustes alignment. Identification accuracy and differential identifiability were calculated for each combination of sessions.

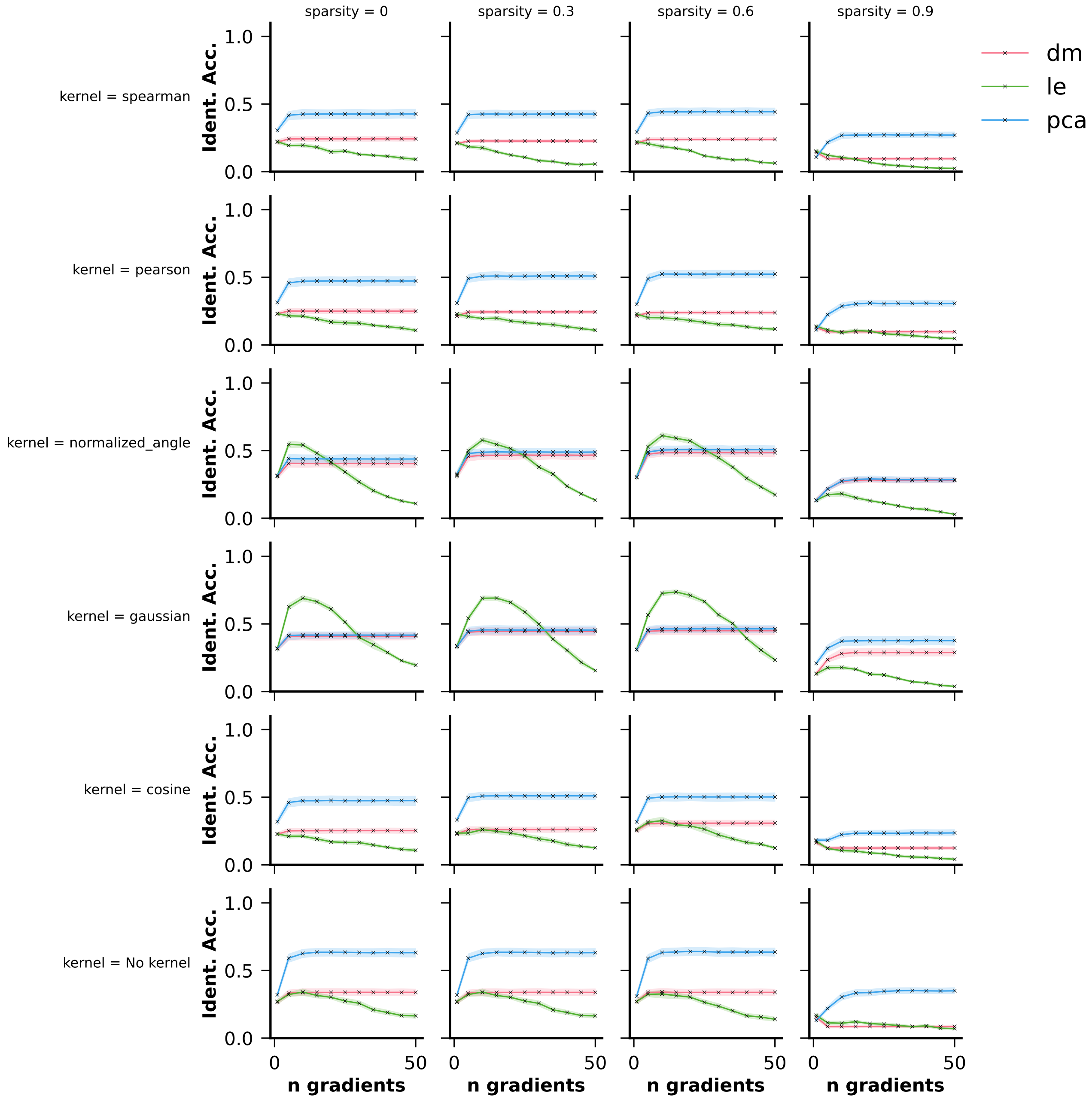


**Figure S2.** Identification accuracy (y-axis) across different kernels (rows), sparsities (columns), and dimensionality reduction approaches (hue) for varying numbers of gradients (x-axis) used in Procrustes alignment. FC gradients were extracted using the Schaefer 100 parcellation.

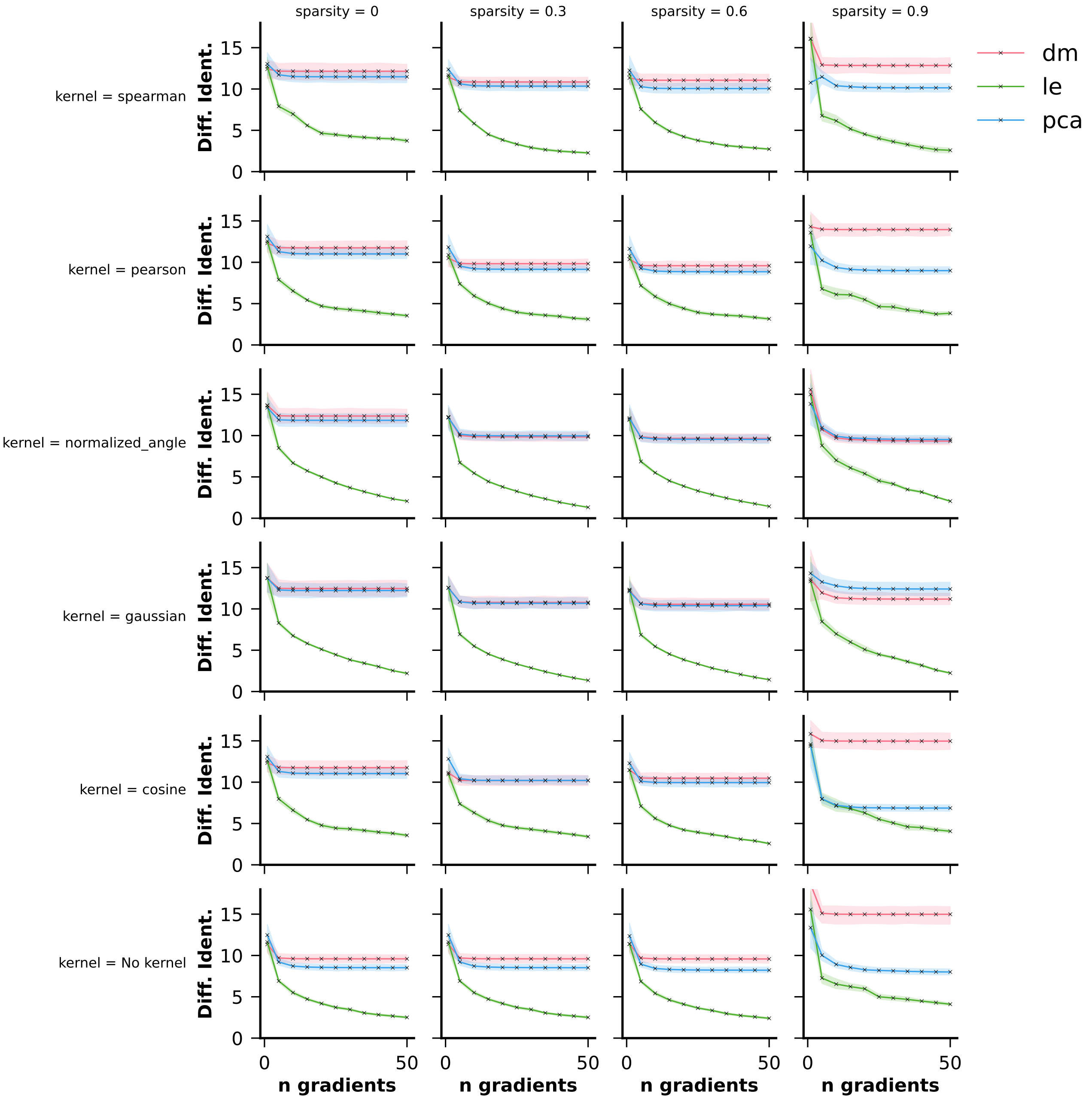


**Figure S3.** Differential identifiability (y-axis) across different kernels (rows), sparsities (columns), and dimensionality reduction approaches (hue) for varying numbers of gradients (x-axis) used in Procrustes alignment. FC gradients were extracted using the Schaefer 100 parcellation.

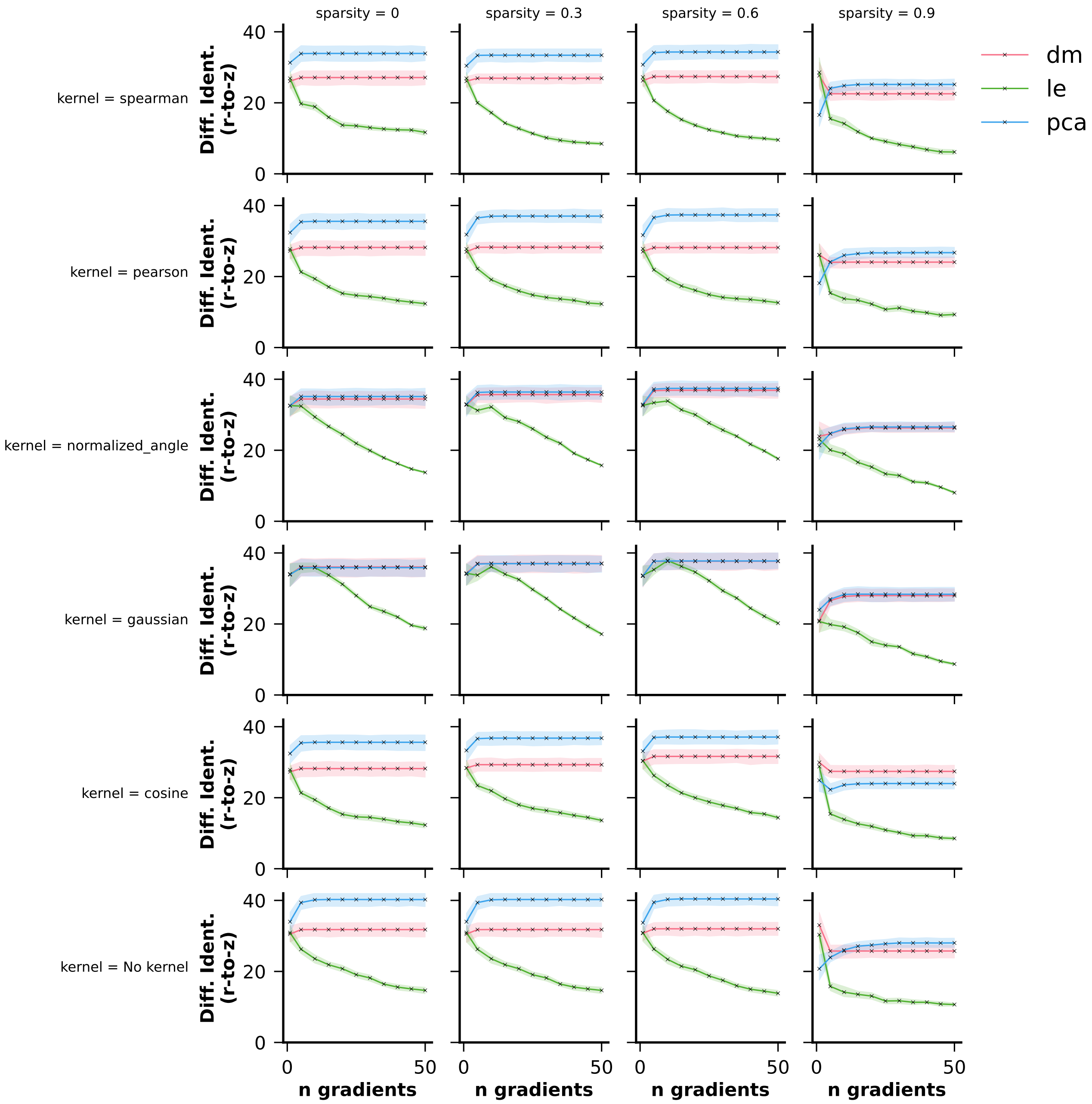


**Figure S4.** Differential identifiability (y-axis) after correlation values underwent Fisher's r-to-z transformation across different kernels (rows), sparsities (columns), and dimensionality reduction approaches (hue) for varying numbers of gradients (x-axis) used in Procrustes alignment. FC gradients were extracted using the Schaefer 100 parcellation.

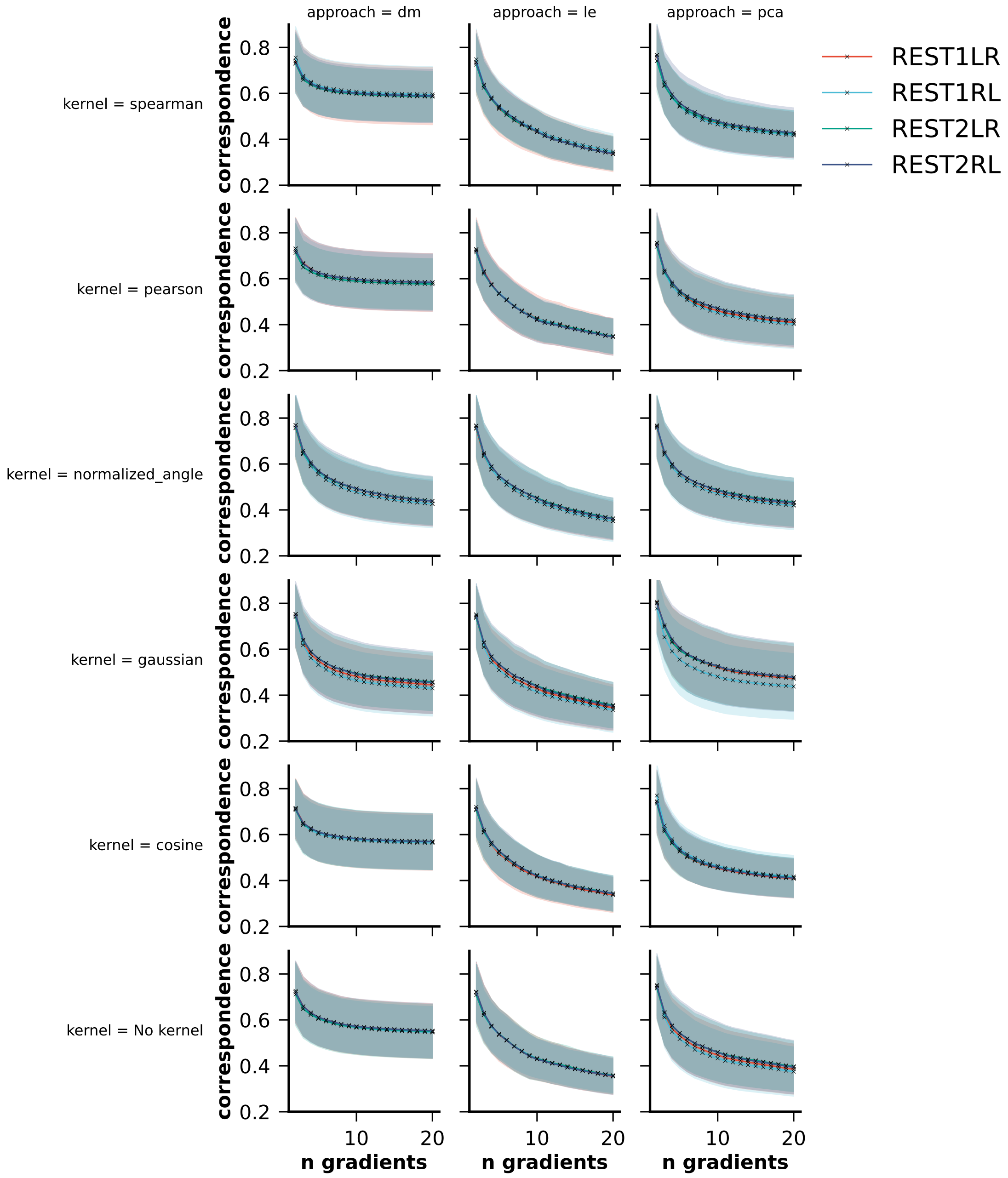


**Figure S5.** The correspondence between the unaligned and the aligned principal gradient per subject per session calculated using the transformation matrices. FC gradients were extracted using the Schaefer 100 parcellation and different kernels (rows) as well as dimensionality reduction approaches (columns).

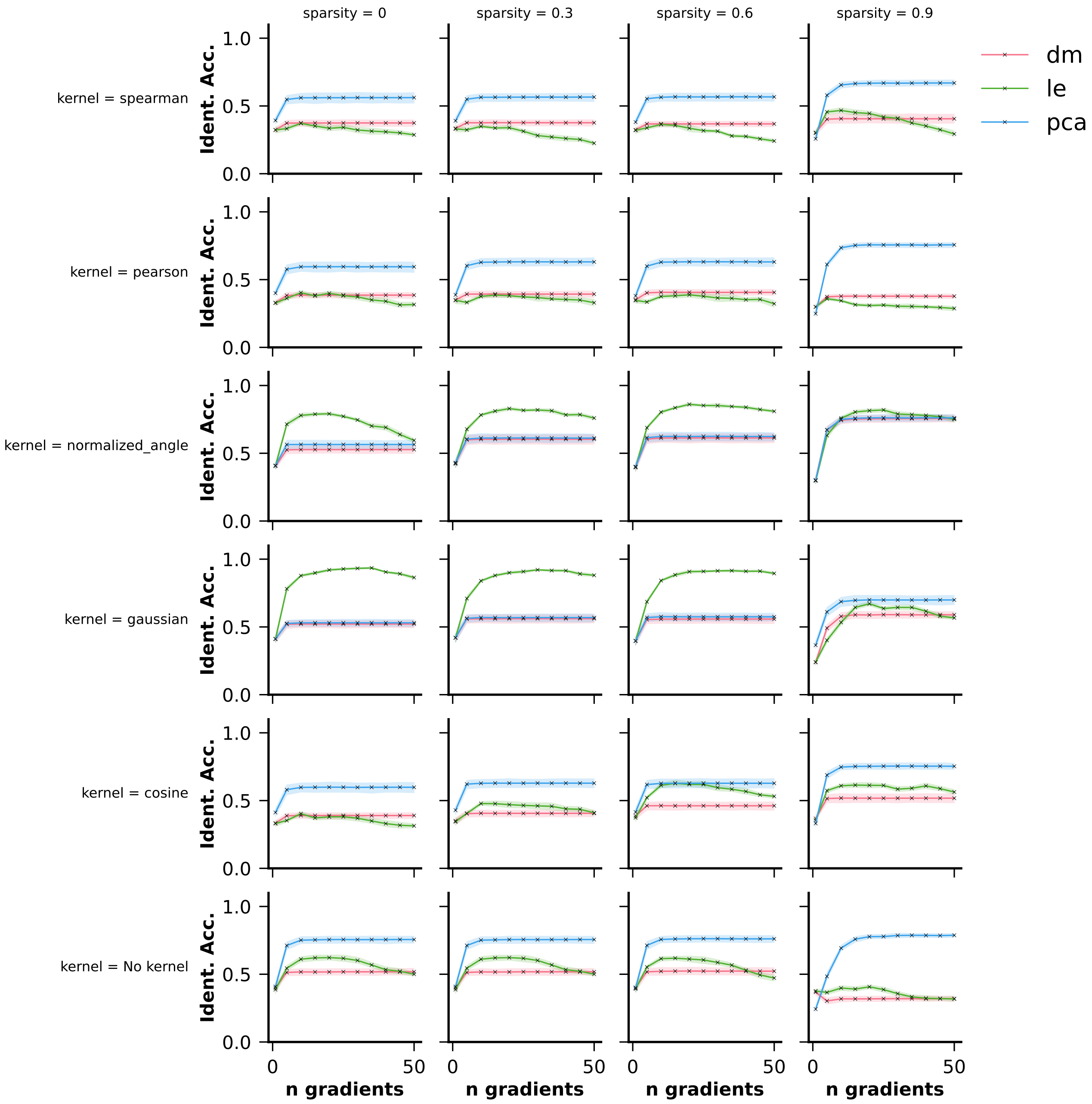


**Figure S6.** Identification accuracy (y-axis) across different kernels (rows), sparsities (columns), and dimensionality reduction approaches (hue) for varying numbers of gradients (x-axis) used in Procrustes alignment. FC gradients were extracted using the Schaefer 200 parcellation.

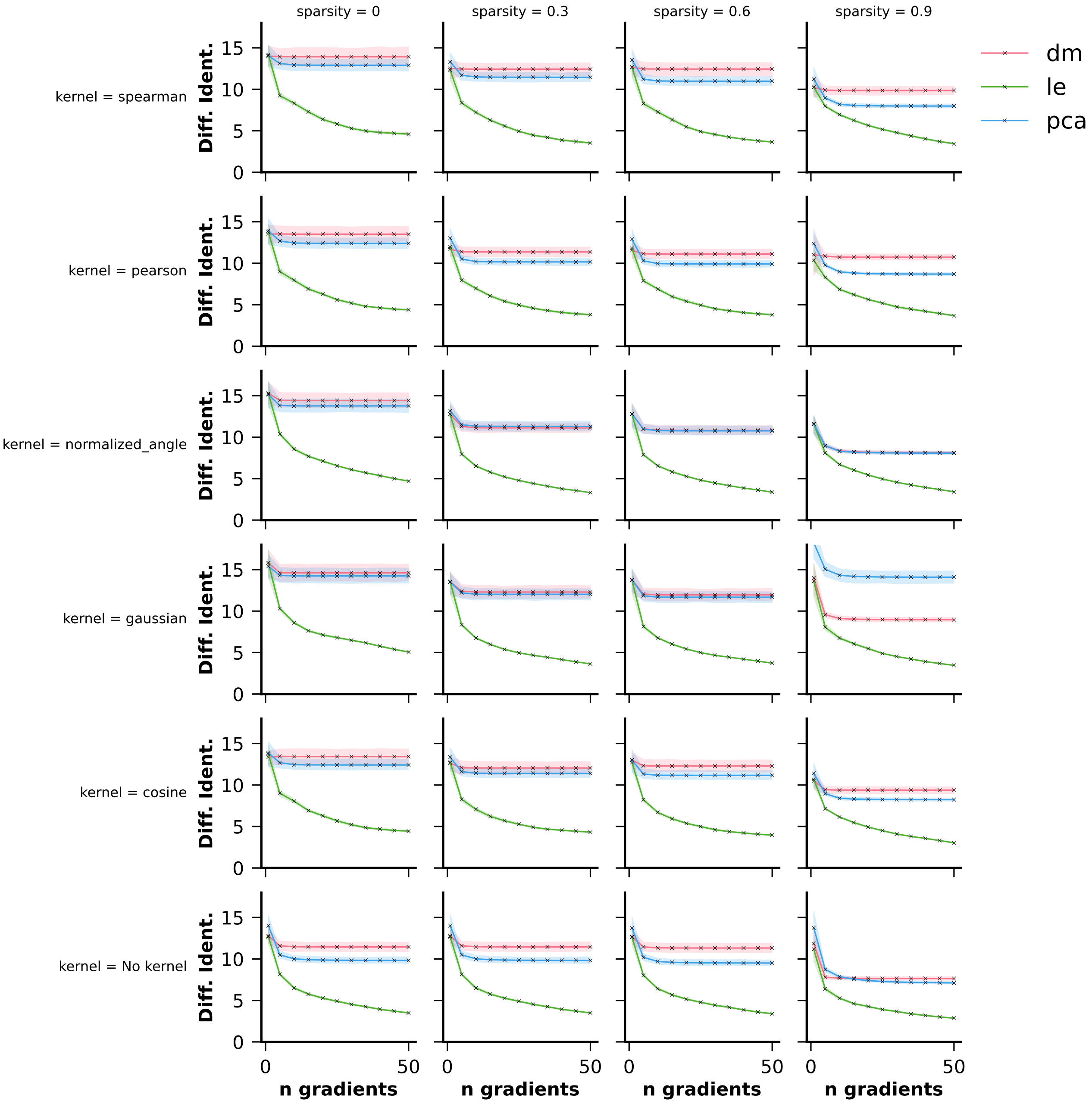


**Figure S7.** Differential identifiability (y-axis) across different kernels (rows), sparsities (columns), and dimensionality reduction approaches (hue) for varying numbers of gradients (x-axis) used in Procrustes alignment. FC gradients were extracted using the Schaefer 200 parcellation.

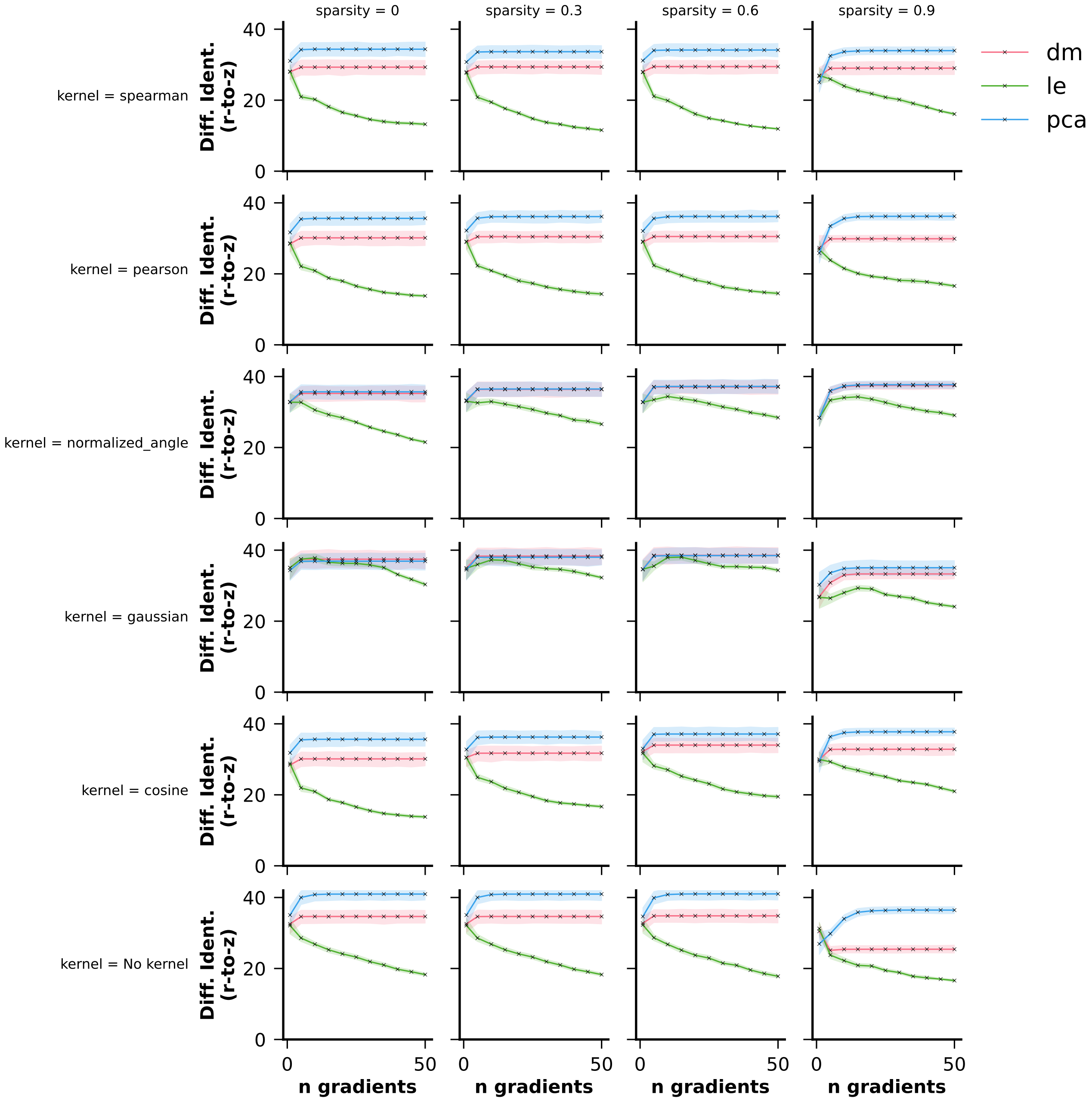


**Figure S8.** Differential identifiability (y-axis) after correlation values underwent Fisher's r-to-z transformation across different kernels (rows), sparsities (columns), and dimensionality reduction approaches (hue) for varying numbers of gradients (x-axis) used in Procrustes alignment. FC gradients were extracted using the Schaefer 200 parcellation.

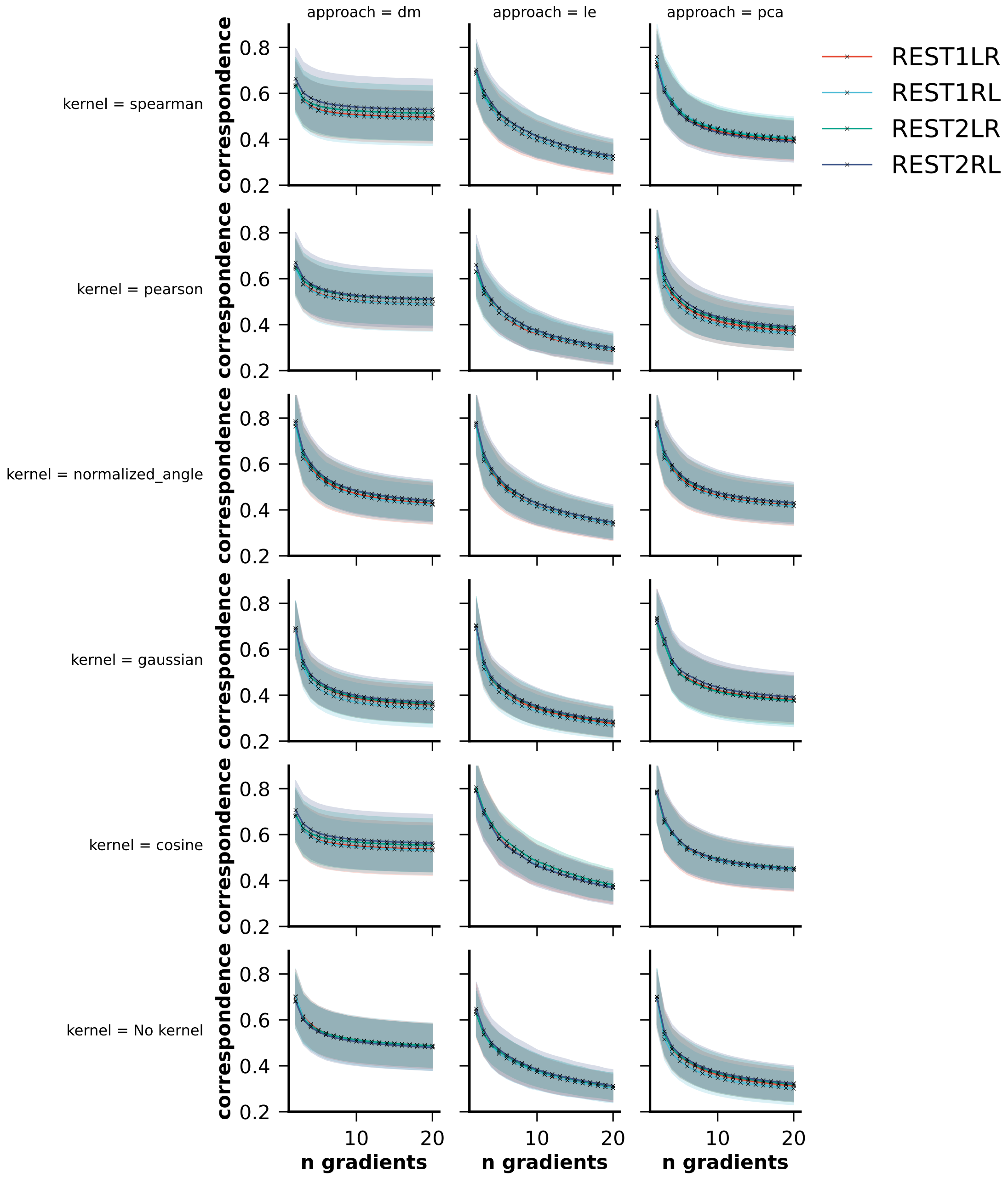


**Figure S9.** The correspondence between the unaligned and the aligned principal gradient per subject per session calculated using the transformation matrices. FC gradients were extracted using the Schaefer 200 parcellation and different kernels (rows) as well as dimensionality reduction approaches (columns).

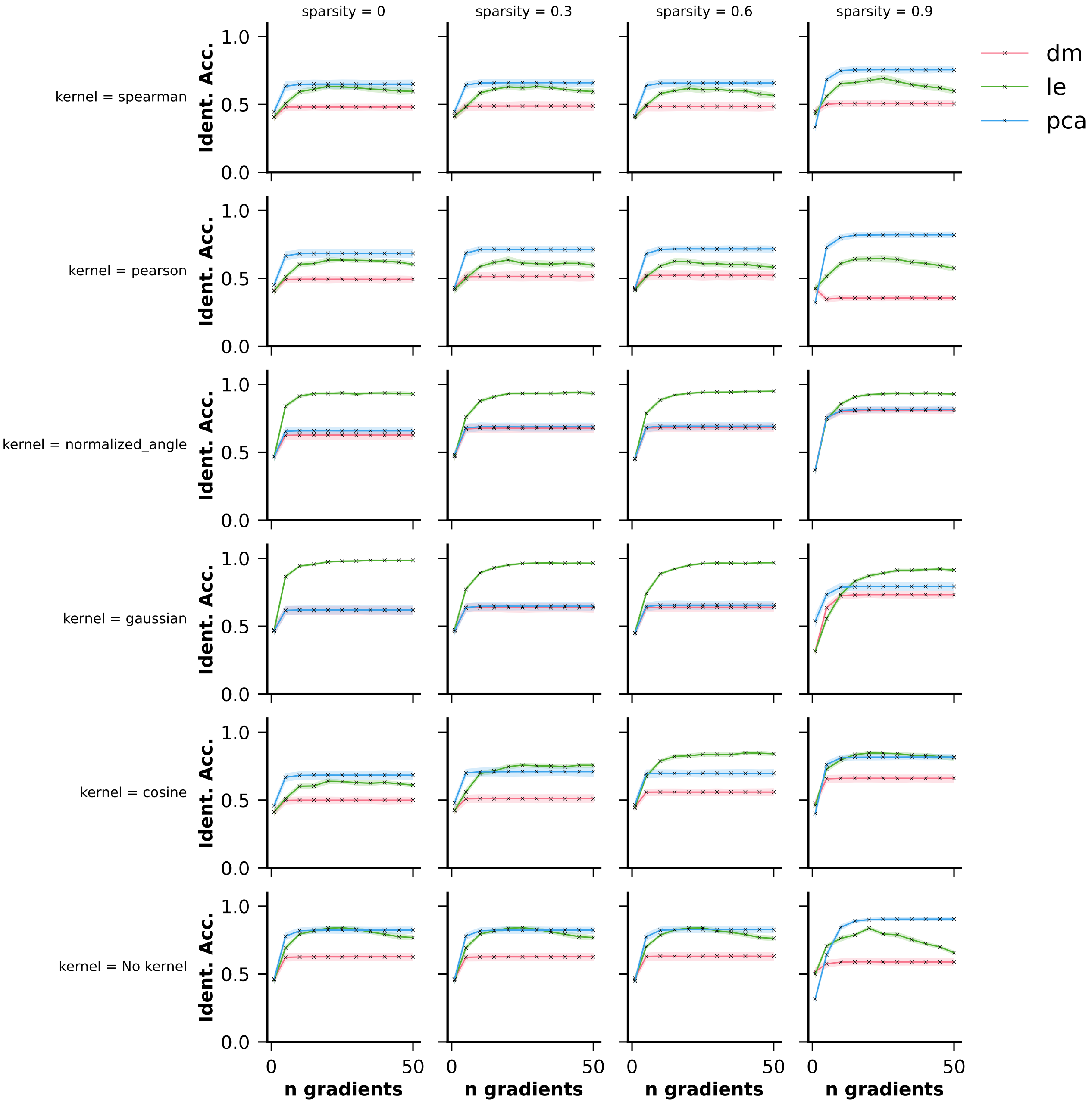


**Figure S10.** Identification accuracy (y-axis) across different kernels (rows), sparsities (columns), and dimensionality reduction approaches (hue) for varying numbers of gradients (x-axis) used in Procrustes alignment. FC gradients were extracted using the Schaefer 400 parcellation.

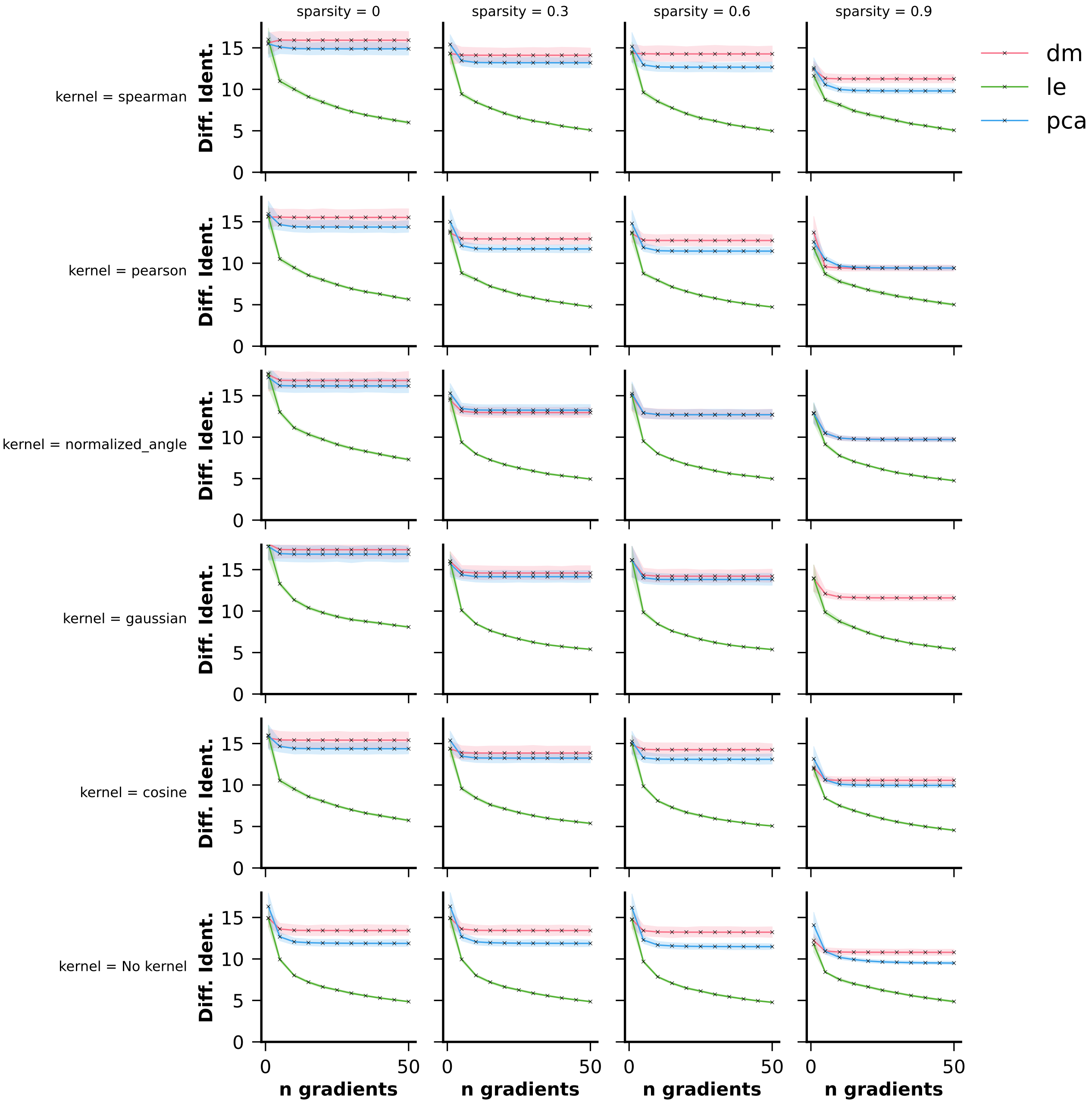


**Figure S11.** Differential identifiability (y-axis) across different kernels (rows), sparsities (columns), and dimensionality reduction approaches (hue) for varying numbers of gradients (x-axis) used in Procrustes alignment. FC gradients were extracted using the Schaefer 400 parcellation.

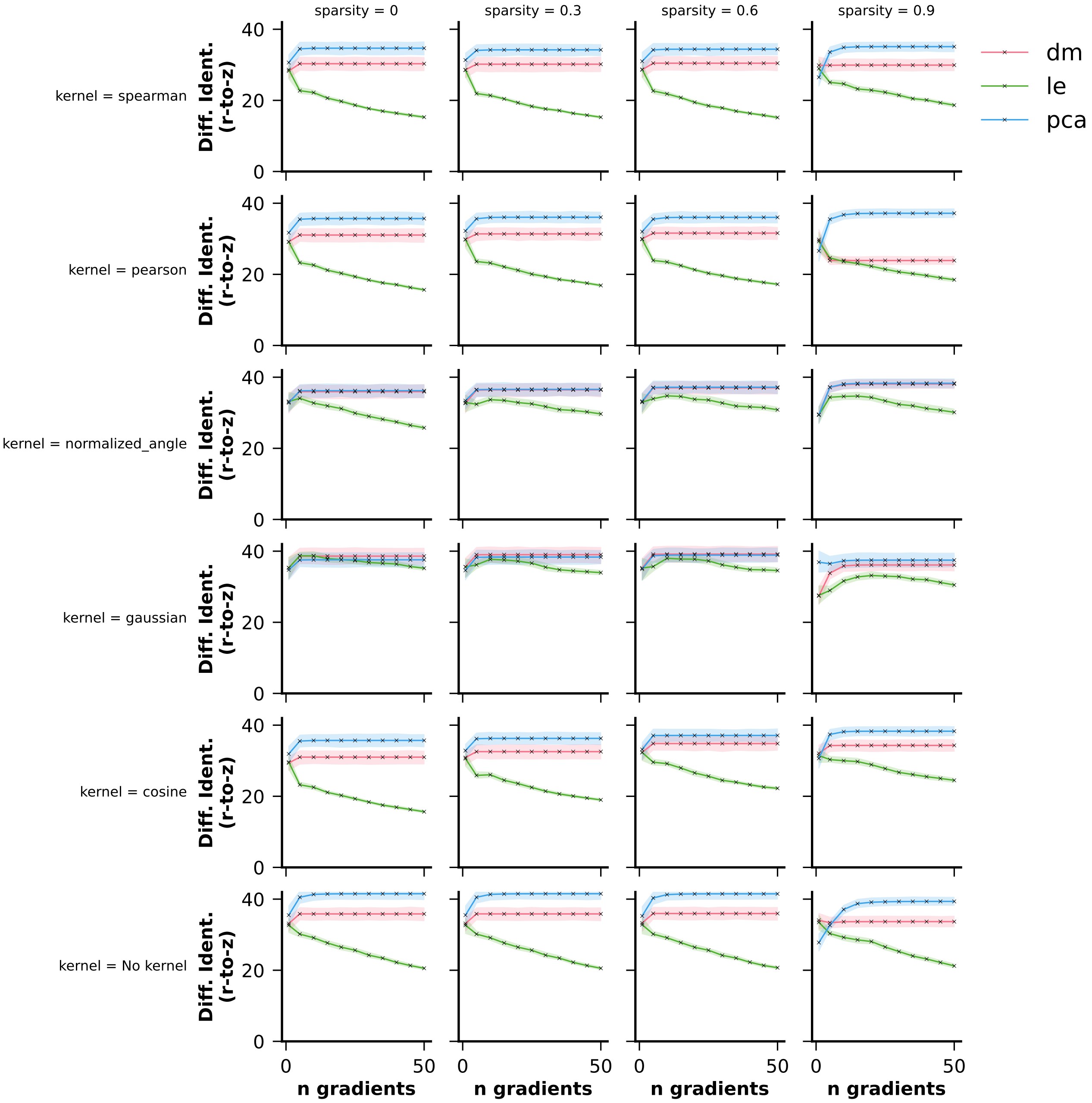


**Figure S12.** Differential identifiability (y-axis) after correlation values underwent Fisher's r-to-z transformation across different kernels (rows), sparsities (columns), and dimensionality reduction approaches (hue) for varying numbers of gradients (x-axis) used in Procrustes alignment. FC gradients were extracted using the Schaefer 400 parcellation.

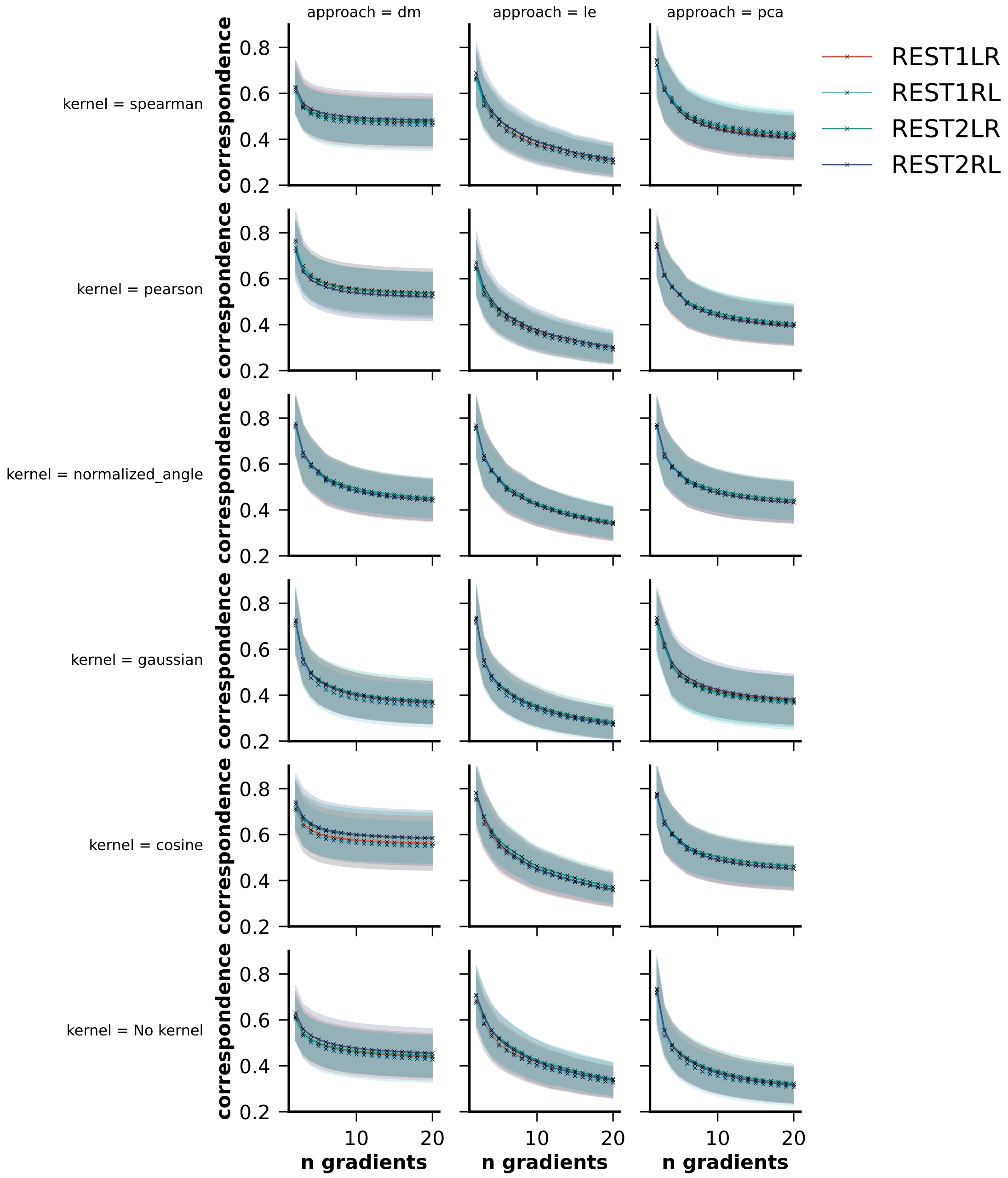


**Figure S13.** The correspondence between the unaligned and the aligned principal gradient per subject per session calculated using the transformation matrices. FC gradients were extracted using the Schaefer 400 parcellation and different kernels (rows) as well as dimensionality reduction approaches (columns).

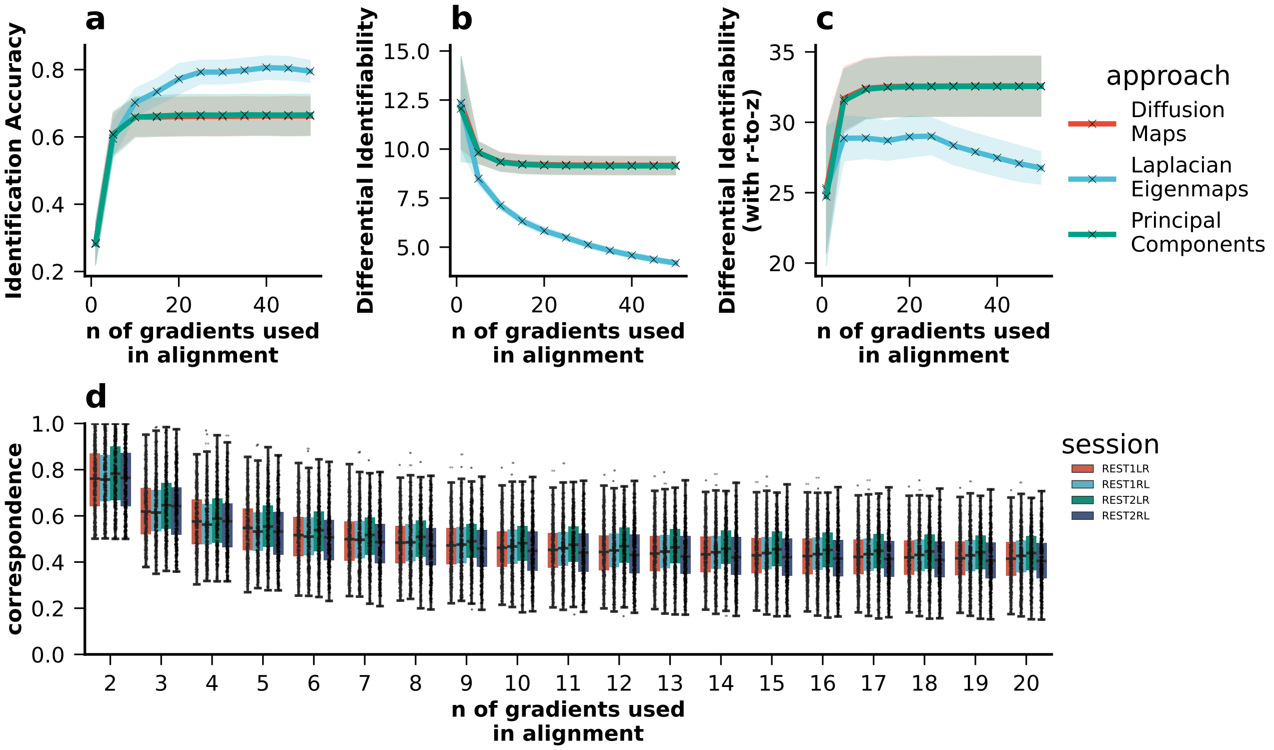


**Figure S14.** Impact of Procrustes alignment on **a)** identification accuracy and **b)** differential identifiability in the HCP-YA dataset without using ICA-FIX motion correction or motion regression. For each subject, gradients were extracted per session (kernel = normalized_angle; sparsity = 0.9). They were then aligned to the holdout reference gradient using Procrustes alignment. Identification accuracy and differential identifiability were calculated for each combination of sessions (NSessions= 4; NCombinations=6). **c)** Lastly, the correspondence between the unaligned and the aligned principal gradient per subject per session were calculated using the transformation matrices.

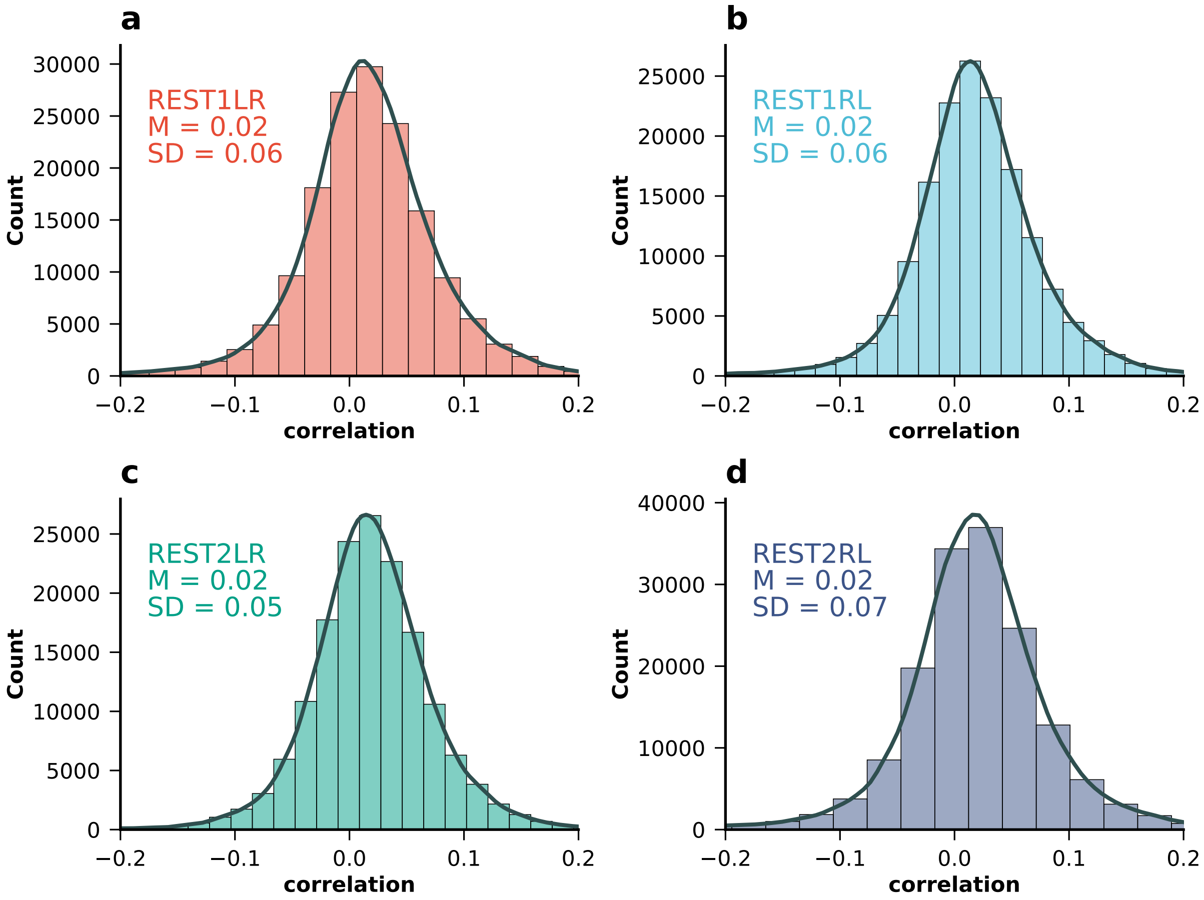


**Figure S15.** Distribution of correlations between each subjects' ROI time series and FD time series for all four resting state fMRI sessions in the HCP-YA dataset without using ICA-FIX motion correction or motion regression: **a)** REST1LR, **b)** REST1RL, **c)** REST2LR, and **d)** REST2RL.

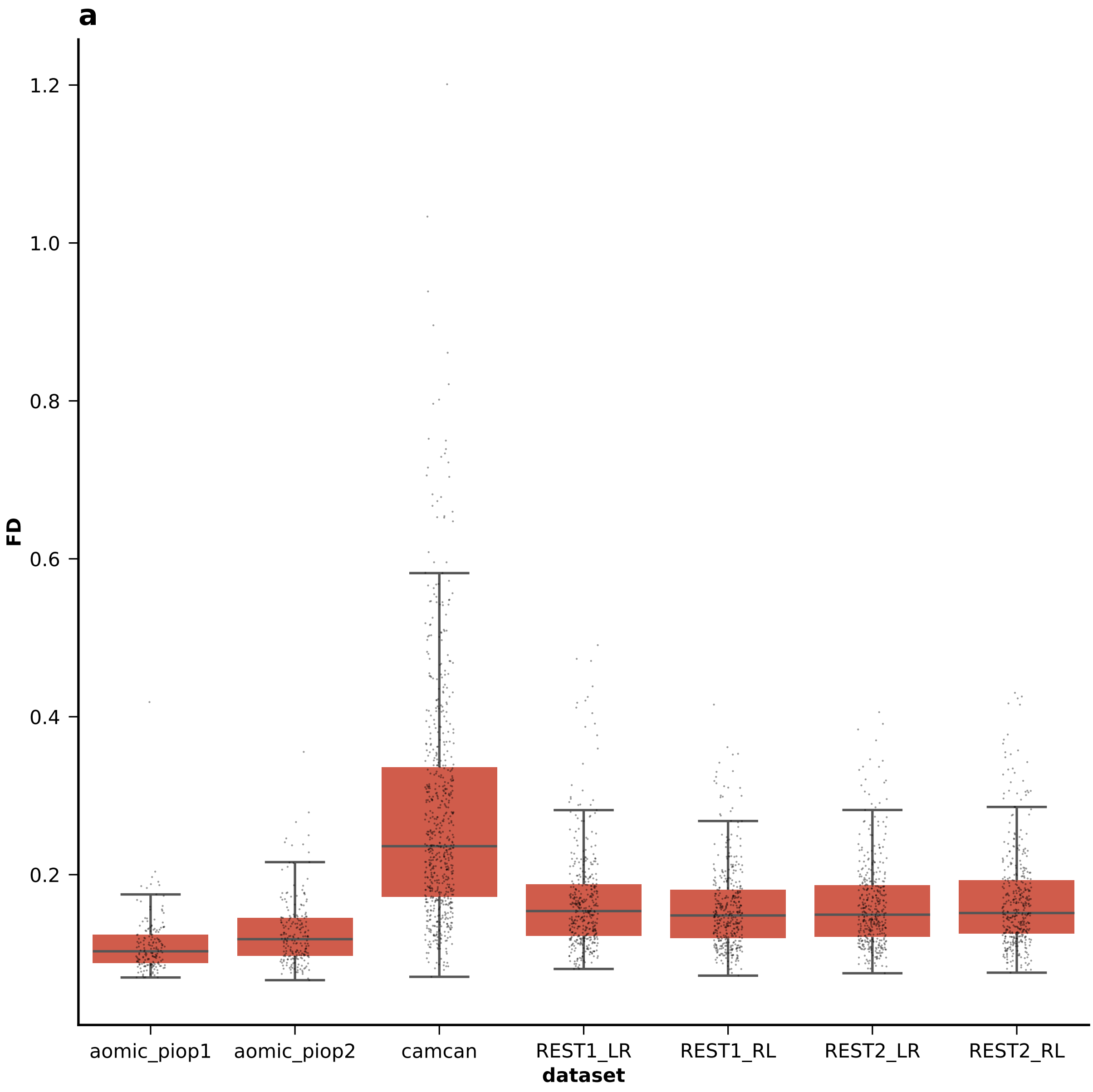


**Figure S16.** Distribution of FD values for all datasets
